# Optimal spatial allocation of enzymes as an investment problem

**DOI:** 10.1101/2021.12.16.473072

**Authors:** Giovanni Giunta, Filipe Tostevin, Sorin Tănase-Nicola, Ulrich Gerland

**Affiliations:** Physics of Complex Biosystems, Physik-Department, Technische Universität München, James-Franck-Str. 1, 85748 Garching, Germany; Department of Cell and Molecular Biology, Uppsala University, Sweden

## Abstract

Given a limited number of molecular components, cells face various allocation problems demanding decisions on how to distribute their resources. For instance, cells decide which enzymes to produce at what quantity, but also where to position them. Here we focus on the spatial allocation problem of how to distribute enzymes such as to maximize the total reaction flux produced by them in a system with given geometry and boundary conditions. So far, such distributions have been studied by computational optimization, but a deeper theoretical understanding was lacking. We derive an optimal allocation principle, which demands that the available enzymes are distributed such that the marginal flux returns at each occupied position are equal. This ‘homogeneous marginal returns criterion’ (HMR criterion) corresponds to a portfolio optimization criterion in a scenario where each investment globally feeds back onto all payoffs. The HMR criterion allows us to analytically understand and characterize a localization-delocalization transition in the optimal enzyme distribution that was previously observed numerically. In particular, our analysis reveals the generality of the transition, and produces a practical test for the optimality of enzyme localization by comparing the reaction flux to the influx of substrate. Based on these results, we devise an additive construction algorithm, which builds up optimal enzyme arrangements systematically rather than by trial and error. Taken together, our results reveal a common principle in allocation problems from biology and economics, which can also serve as a design principle for synthetic biomolecular systems.

## INTRODUCTION

Living systems often need to allocate limited resources to different tasks. For instance, this situation arises when a colony of microbes differentiates into different cell types, each specialized to a task that contributes to the growth of the colony as a whole [1]. Cells solve this allocation problem by regulating the splitting ratios between cell types during differentiation. Conceptually similar allocation problems arise on the molecular level, where enzymes need to be allocated to different targets. For instance, a limited number of ribosomes must produce different types of proteins in different ratios to achieve balanced cell growth [2–4]. Accordingly, cells dynamically control these ratios when their environment changes to support different growth rates [5, 6]. Cells also control the spatial localization of enzymes to optimize or regulate metabolic fluxes [7, 8]. These and many other biological examples share the basic characteristics of generic resource allocation problems, in which a given amount of a resource must be distributed among competing alternatives to maximize the expected performance of the system [9].

The optimal allocation of limited resources has been extensively studied in economics. Rules such as Kelly’s criterion [10] and returns-variance analysis [11] are used to determine optimal betting strategies or optimal portfolios [12]. These criteria typically assume that the expected return on a bet only depends on its placement and not on the placement of other bets. Although there are biological systems for which such criteria can be directly applied [13], the treatment needs to be generalized to describe systems, for which the performance depends in coupled, nonlinear ways on the amounts of the resource allocated to each task. Here we show that the problem of allocating enzymes to different spatial positions to optimize the total enzymatic reaction flux has a one-to-one mapping to an investment problem where investments and returns are interdependent. In this context, enzyme molecules represent capital that can be invested at different positions within the system. Each position corresponds to an asset generating a certain expected return in terms of reaction flux. However, the placement of enzymes affects the substrate profile, and thereby the flux return from other enzymes. Hence, the spatial enzyme allocation problem corresponds to a betting problem where the coupling between the returns and the bets is determined by the reaction-diffusion dynamics of the substrate.

We consider a general class of models (Fig. 1), in which the substrate 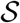 can enter an arbitrarily shaped system at (subsections of) the exterior or interior boundaries, while internal 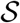 is transported via diffusion and possibly also by advection, and 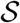 is lost within the system to competing reactions or at the boundaries due to leakage. The enzymes 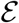 can be freely arranged on the boundaries and within the system. Potential co-localization of 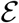 with other enzymes [8, 14–16] is incorporated in our model to the extent that the substrate influx at the boundaries may be caused by other, boundary-localized enzymes.

**Figure 1.**
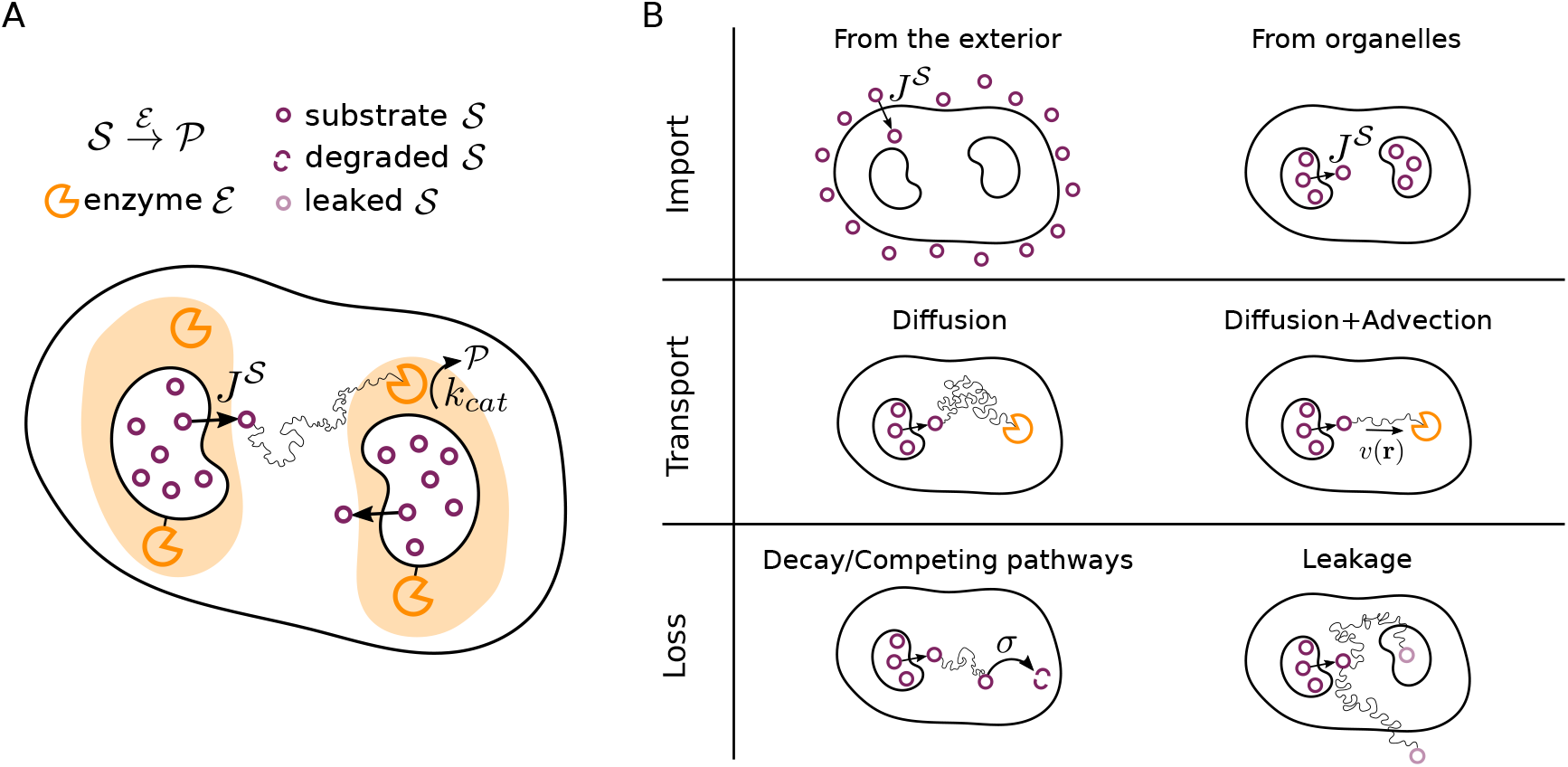
Schematic representation of the class of models considered in this work. (A) An enzyme 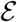 converts a substrate 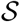 to product 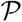 with a catalytic rate *k*_cat_. The enzyme molecules can be freely placed, both within the system and on its interior or exterior membranes. (B-Import) 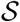 is either imported from the exterior or produced within internal compartments. (B-Transport) Within the system, 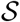 is transported diffusively, and possibly also advectively, e.g. by active transport or cytoplasmic streaming. (B-Loss) During transport, 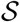 can also be degraded with rate *σ* or leak out of the system via a membrane.

A previous numerical study of a minimal one-dimensional model within the class of models of Fig. 1 observed a localization-delocalization transition in the optimal enzyme distribution as a function of a dimensionless reaction-diffusion parameter [17]: When reactions are slow compared to diffusion, localizing all 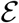 at the source boundary is optimal, while a more extended profile with some enzymes also in the interior is optimal for faster reactions. The existence of this transition was subsequently found to be robust with respect to the spatial dimension and specific loss mechanism and reaction kinetics [18]. However, the physical principles governing the optimal configurations and the generality of the transition have remained elusive, and a construction principle for optimal enzyme arrangements is lacking. Within the economics analogy, the localization-delocalization transition corresponds to a transition in investment strategy from investing everything on a single asset to having a diversified portfolio with multiple assets. Can we explain the generality of the transition by analogy with a diversification investment strategy? Moreover, just as Kelly’s criterion [10] and returns-variance analysis [11] provide a set of rules for the optimal partition of bets, can we derive a generalized criterion to construct optimal enzyme arrangements?

We address these open questions with a variational approach, which ultimately reveals the investment strategy underlying optimal enzyme arrangements. The practical significance of our theoretical analysis has three aspects. First, it produces a criterion to determine, for a given experimental situation, whether enzyme localization is optimal. Second, we turn this criterion into an additive construction scheme for optimal enzyme arrangements. This is relevant for synthetic biology [19], where spatial optimization of enzymatic systems is used to boost yields of useful products such as drug molecules [20–22] and biofuels [23, 24]. Third, the conceptual framework underlying our analysis and the economic interpretation that it provides can be transferred to other biological or economical allocation problems. This supports the agenda of rationalizing biological strategies with economic principles such as bet-hedging [25–27], division of labor [28], and Pareto fronts [29].

## RESULTS

### Variational approach to the spatial enzyme allocation problem

We consider catalytic reactions where a substrate 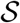 is converted into a product 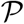 by an enzyme 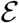 (Fig. 1A). The substrate can enter the system from the exterior or from an internal compartment, e.g. an organelle (Fig. 1B-Import). Enzymes may be located both in the bulk of the sytem and at the boundaries where the substrate enters. The former fraction is described by the concentration field *e*(**r**), with **r** denoting positions inside the system, while *e_S_*(**s**) is the density of 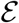 on the boundary surface at position **s**. The concentration of 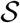 at time *t* is *ρ*(**r**,*t*). We consider the distribution of 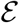 as stationary and explore all possible spatial arrangements of the enzymes. The rate at which enzymes locally catalyze product formation depends on the substrate concentration *ρ*(**r**,*t*). We express this dependence in the general form *k*_cat_*eF*[*ρ*], where *k*_cat_ is the catalytic rate of the enzyme and *F*[*ρ*] is a monotonically-increasing reaction function that depends on the enzyme’s reaction mechanism, e.g., *F*[*ρ*] = *ρ*/(*K*_M_ + *ρ*) for Michaelis-Menten kinetics and *F*[*ρ*] ∝ *ρ* for linear mass-action kinetics.

Within the system, the transport of substrate molecules (Fig. 1B-Transport) can be both stochastic, with a diffusion coefficient *D*, as well as directed, with a velocity field **v**(**r**), where the latter could be due to cytoplasmic streaming [30] or cargo-carrying molecular motors [31]. If the substrate is intrinsically unstable [32] or subject to competing reaction pathways [16], it decays with rate constant *σ* (Fig. 1B-Loss). The dynamical interplay of these processes then follows

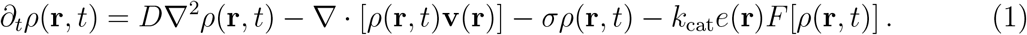

In the following, we focus on steady-state conditions, *∂_t_ρ*(**r**,*t*) = 0, where *ρ*(**r**,*t*) = *ρ*(**r**). We consider two types of boundaries for the system. First, boundaries at which substrate enters. Including reactions of substrate with enzymes that may be located at such a boundary, we have a boundary condition of the form

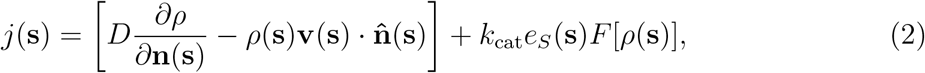

where *j*(**s**) is the influx of 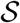 at position **s**, 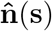 is a unit vector normal to the boundary, pointing outwards from the system, and 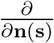 represents the magnitude of the normal component of the gradient. This condition enforces flux conservation at the boundary: the influx *j*(**s**) must equal the sum of the local transport and reaction flux. We allow for *j*(**s**) to be a discontinuous function of **s** as there could be regions on the boundary without influx, *j*(**s**) = 0 (see Appendix A for details on how such discontinuities are treated). The second kind of boundaries we consider are boundaries at which substrate is lost (Fig. 1B-Loss), e.g. via leakage through a membrane [33]. For these, we impose absorbing boundary conditions, *ρ*(**s**) = 0. For convenience, we represent these boundary conditions as

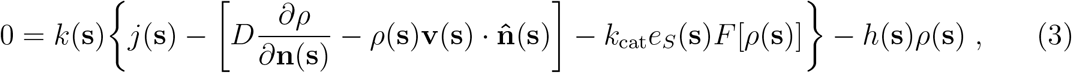

where *k*(**s**) and *h*(**s**) are set to represent absorbing boundaries (*k* = 0, *h* =1) or boundaries with influx of substrate (*k* = 1, *h* = 0).

The interplay of processes of the type illustrated in Fig. 1 and described by Eqs. 1 to 3 have been demonstrated to generate stable gradients of intracellular components in a variety of biological contexts [34–37]. An immediate consequence of non-uniform substrate profiles is that different spatial enzyme distributions *e*(**r**) generate different total reaction fluxes. In cases where the substrate is produced inside organelles, e.g. mitochondria or endoplasmatic reticula, the associated enzymes tend to be concentrated in proximity of the organelle membrane but also present in the cytoplasm [38–41]. In contrast, when the substrate is imported into the cytoplasm from the external environment, some of the associated metabolic enzymes are localized to the external membrane [42].

The total reaction flux 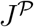, i.e. the rate at which the whole system produces 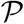, has contributions from enzymes at the boundaries and within the bulk,

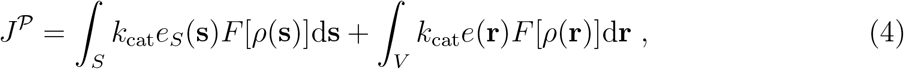

where the first integral is over all boundary surfaces *S* and the second is over the volume *V* of the system. Given a fixed amount of available enzymes, *E_T_* = *∫_S_ e_S_*(**s**)d**s** + *∫_v_ e*(**r**)d**r**, how should these enzymes be positioned such as to maximize the rate 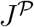 of product formation?

This optimization problem can be approached by defining a functional of the form 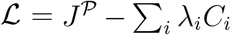, where *C_i_* represent different constraints with associated Lagrange multipliers *λ_i_*. For our model, these constraints are that (i) the total amount of enzymes must equal *E_T_*, (ii) *ρ*(**r**) and *e*(**r**) must jointly satisfy the reaction-diffusion-advection equation, Eq. 1, at each point **r**, and (iii) *ρ*(**s**) and *e_S_*(**s**) must jointly satisfy the boundary condition, Eq. 3, at each boundary point **s**. The resulting Lagrangian has the form

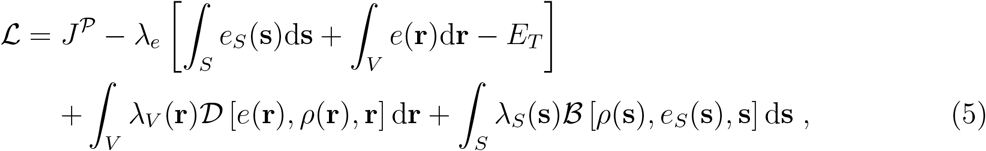

where 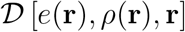 and 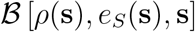 represent the right-hand side of Eqs. 1 and 3 respectively, and the signs in front of the Lagrange multipliers have been chosen such that they result to be positive. Maximizing 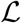 with respect to *e_S_*(**s**) and *e*(**r**) while simultaneously minimizing with respect to all Lagrange multipliers yields the optimal enzyme arrangements satisfying the constraints.

The optimal enzyme arrangement may consist of both regions with a finite density of enzymes and regions with no enzymes. Indeed, previous numerical studies have shown that the optimal enzyme distributions are often discontinuous [16–18]. To account for such discontinuities, we split the volume *V* into different sub-spaces (and similarly for the boundaries *S*); within each sub-space the enzyme density must be continuous and *e*(**r**) ≥ 0, but at the interfaces continuity is not required. The number of sub-spaces as well as the shape of the interface between each pair of sub-spaces are then also variables to be optimized (see Appendix A for details and Appendix B, E, F for examples). This is equivalent to applying the Karush-Kuhn-Tucker conditions [43, 44] on the Lagrangian, Eq. 5, while imposing *e*(**r**) ≥ 0.

We will see below that the variational approach is useful even when it is impossible to solve for the optimal enzyme arrangement analytically. In cases where it is possible to extract the exact functional form of the optimal profile, the problem can usually be reduced to a low-dimensional optimization over a small number of parameters, which is then easily performed numerically (see Appendix E for an example). For systems where the functional form of *e*(**r**) cannot be found analytically, such as those with complex geometries, we will describe a scheme for constructing the optimal enzyme arrangement.

### Enzyme allocation problem as a betting game

The spatial enzyme allocation problem defined above can be mapped to a betting game. We consider a game that has a single winning outcome out of a set of mutually exclusive events. Each outcome *i* has a certain probability *pi* of being the winning event. We assume that a gambler bets a total amount *B* = ∑_*i*_ *b_i_* with *b_i_* ≥ 0 denoting the bet on outcome *i*. If *i* is the winning event, the gambler wins the amount *α_i_b_i_*, with *α_i_* ≥ 0 referred to as the odds of event *i*. The expected amount of capital won by the gambler is thus *C* = ∑_*i*_ *α_i_b_i_p_i_*. If the expected odds *α_i_p_i_* are independent of the bets *b_i_*, the optimal strategy for a given total budget *B* is to invest everything in the outcome with highest expected odds. However, if the expected odds are coupled to the bets, the optimization of *C* subject to the budget constraint becomes nontrivial, and diversification of bets may be the optimal strategy.

By comparing the objective function *C* of the betting game to the reaction flux 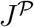 of our spatial enzyme allocation problem, we see that the set of events {*i*} maps to the set of possible enzyme positions, a bet *b_i_* corresponds to the enzyme concentration *e*(**r**) allocated to **r**, and the fixed budget *B* corresponds to the total enzyme amount *E_T_*. Moreover, the probability of *i* being the winning event maps to the probability density for a substrate molecule to bind to an enzyme at position **r**, *p_i_* ↔ *F*[*ρ*(**r**)], and the odds correspond to the catalytic rate *k*_cat_.

In the enzyme system, part of the substrate is lost via the mechanisms of Fig. 1B-Loss, which can be included in the betting game via events with finite probability but zero odds. However, the objective function *C* = ∑_*i*_ *α_i_b_i_p_i_* needs to be generalized to account for the nonlinearity and spatial coupling in the enzyme system: The “expected odds” *k*_cat_*F*[*ρ*(**r**)] depend on the “bets” *e*(**r**), since the binding between enzymes and substrate at a given location generates a diminishing returns effect, i.e., the addition of extra enzymes at that position becomes gradually less efficient. Moreover, the substrate concentrations at different positions are coupled, due to the substrate diffusion (and possibly advection), such that the substrate concentration at one position ultimately depends on the enzymes at every position. This means that the bets placed on one event *i* also affect the winning probabilities *p_j_* with *j* ≠ *i*. These effects are captured by making the probabilities *p_i_* a function of the vector of bets *p_i_* = *f_i_*(**b**) and adding the feedback from the bets onto the winning probabilities via constraints in the objective function,

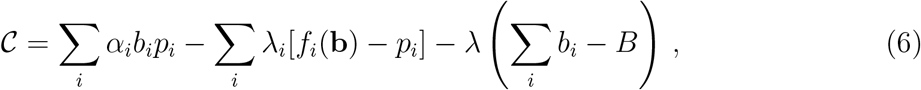

where λ_*i*_ are the Lagrange multipliers for the feedback constraint, analogous to λ(**r**) in the Lagrangian 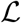 of the enzyme system, Eq. 5. The behavior of this generalized betting game depends on the form of *f_i_*(**b**). In Appendix H, we show for example that in a simple game with two possible winning outcomes, a diminishing returns effect (*∂f_i_*/*∂b_i_* < 0) can result in an optimal strategy with diversified bets (*b*_1_, *b*_2_ > 0).

The above mapping illustrates that in order to solve our optimal enzyme allocation problem, we need to generalize the treatment of betting games to include the coupling between bets and expected gains. Note that we considered only a single round of the betting game so far. Classical treatments of betting games, e.g. the one by Kelly [10], often assume that the gambler iteratively reinvests a certain fraction of the current capital, such that the long-term capital growth rate must be optimized rather than *C*. In the enzyme context, this would correspond to a situation where the product of the enzymatic reaction is used to generate additional enzymes to be placed in the system. Here, we first treat the stationary enzyme allocation problem, corresponding to a repetitive game in which the gambler bets the same total amount *B* in every round, such that the optimal strategy is the same as the optimal strategy for a single round. We will later see that our scheme to construct optimal strategies can also be adapted to the case where a constant fraction of the capital is iteratively reinvested (see below and Appendix H).

### Optimal allocation principle

We now leverage the above variational approach to solve the enzyme allocation problem. Towards this end, we examine the variation of the Lagrangian, Eq. 5, with respect to the enzyme density,

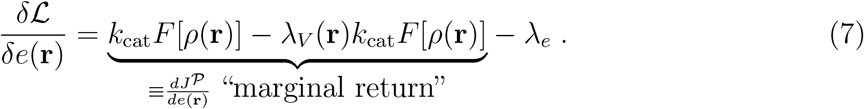

This variation corresponds to the change in the constrained flux 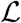 as enzymes are added at position **r**. Three different terms contribute to this change. The first is the increase in flux that would be observed upon adding enzymes at position **r** if everything else in the system were to remain unaffected, which corresponds mathematically to 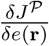. However, changing *e*(**r**) also alters the substrate density profile *ρ*(**r**), thereby affecting the rate of reactions at all positions. This coupling is captured by the second term in Eq. 7, where the Lagrange multiplier λ_*V*_(**r**) ensures that *ρ*(**r**) and *e*(**r**) satisfy the constraint of the reaction-diffusion equation. Thus the sum of the first two terms is the total rate of change of the reaction flux as extra enzymes are added at position **r**. In the following, we refer to this quantity as the “marginal return” on an investment of enzymes, and denote it by 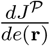. Finally, the third term, 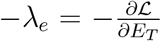, corresponds to a marginal cost of adding extra enzymes into the system. This marginal cost can be interpreted as the reduction in flux from other positions **r**′ ≠ **r** in the optimal configuration, as enzymes are moved from these positions to **r** in order to satisfy the constraint of constant total enzyme number.

In the optimal enzyme profile *e*\*(**r**), we must distinguish between regions where placing enzymes is optimal (*e*\*(**r**) finite), and empty regions where placing enzymes is suboptimal (*e*\*(**r**) = 0). Wherever *e*\*(**r**) > 0, the variation 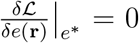. The functional derivative with respect to the surface density 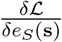 has the same form as Eq. 7. Thus the optimal profile follows a ‘homogeneous marginal returns’ criterion (HMR criterion): it is such that at any position with enzymes, the marginal return on placed enzymes equals a constant value, namely 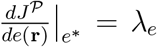. By contrast, in empty regions, the marginal flux gain from placing enzymes is less than the associated marginal cost, 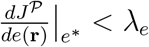.

To illustrate this optimal allocation principle, we apply the HMR criterion to the simple case of a linear reaction (*F*[*ρ*] = *ρ*/*K*_M_) with no advection or decay (|**v**| = *σ* = 0) in a one-dimensional domain of size *L*, with a source of substrate at the left boundary (*x* = 0) and an absorbing boundary on the right (Fig. 2A). This case is exactly solvable within the variational approach, such that we can explicitly see how variations of the enzyme profile affect the marginal returns landscape. Moreover, this case allows us to cross-validate our variational approach with a previous numerical study [17]. As shown in Appendix B, the optimal enzyme profile for this example is

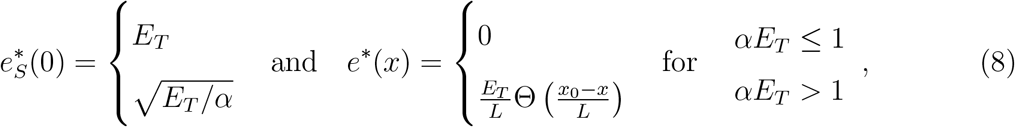

with *α* = *Lk*_cat_/(*K*_M_*D*), the Heaviside step function Θ(*x*), and *x*_0_ = *L* 1 – (*αE_T_*)^−1/2^ Eq. 8 shows that the optimal enzyme profile undergoes a localization-delocalization transition from having all enzymes localized at the source, to an extended profile with enzymes also in the interior of the system (Fig. 2B, orange lines). The transition occurs as a function of the dimensionless parameter *αE_T_*, with *αE_T_* = 1 marking the transition point.

**Figure 2.**
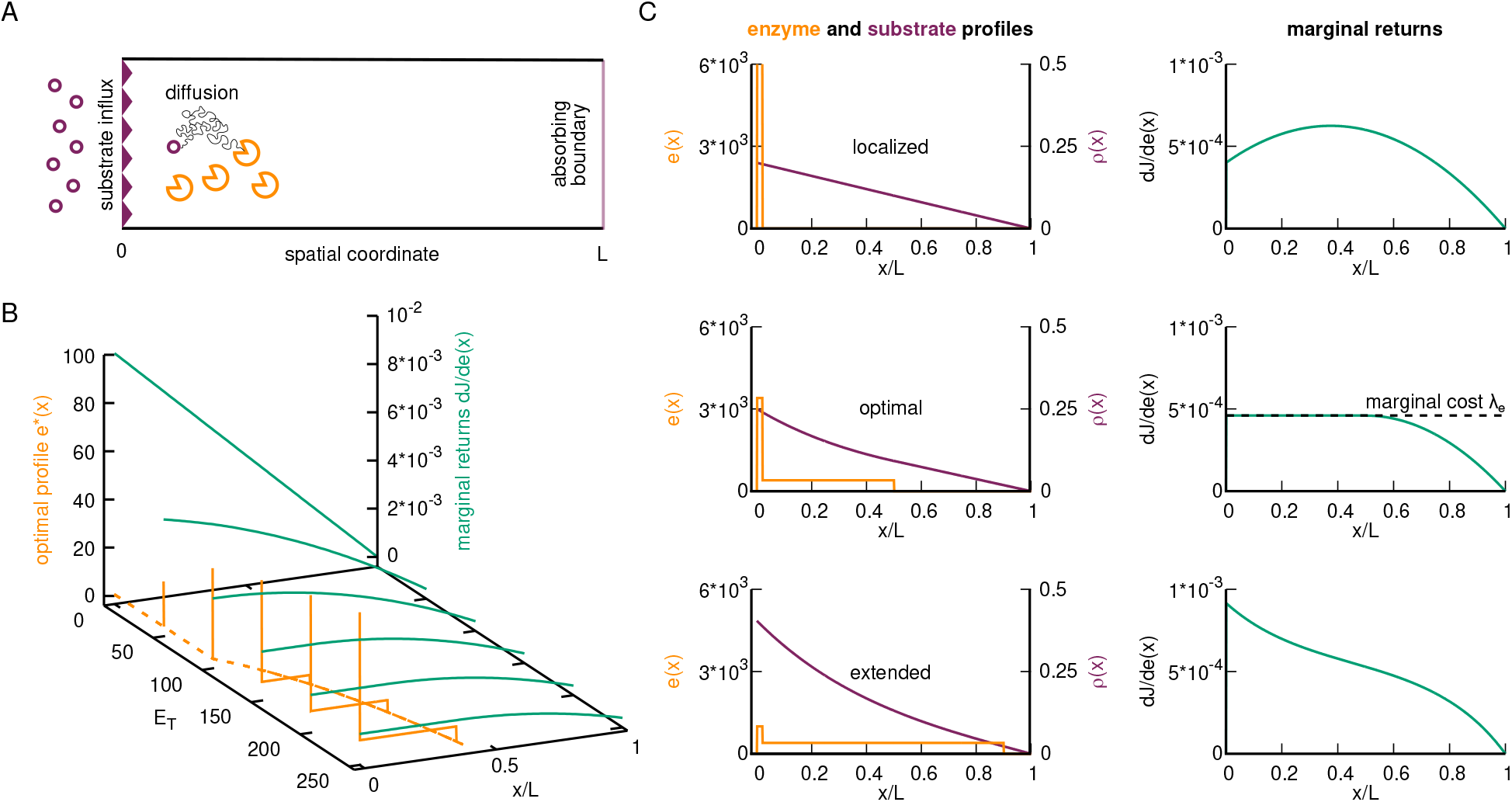
Homogeneous Marginal Returns (HMR) criterion for the optimal spatial allocation of enzymes. (A) Illustration of the analytically solvable 1D model with a source of substrate at *x* = 0 and an absorbing boundary at *x* = *L*. Within the system, the substrate diffuses and can react with the spatially arranged enzymes. (B) Optimal enzyme profile with corresponding marginal return landscape as a function of the total enzyme amounts *E_T_* (the reaction-diffusion parameter is fixed to *α* = 10^-2^). (C) Different enzyme and substrate profiles (left) with corresponding marginal returns (right) for *E_T_* = 400. Top: Localized configuration. The marginal returns landscape is peaked at *x* > 0, indicating that enzymes should be moved away from the source. Middle: Optimal profile with constant marginal return over the region with enzymes (HMR criterion). In the constant region, the marginal return is equal to the marginal cost λ_*e*_ of adding extra enzymes into the system. Bottom: Overextended profile. Monotonically decreasing marginal return, indicating that enzymes should be moved towards the source at *x* = 0.

The analytic solution demonstrates how the optimal enzyme profile arises from a balance between gain and cost. The shape of the marginal returns landscape (Fig. 2B, green lines) changes as the total enzyme amount *E_T_* is increased. Below the transition, the marginal returns 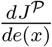 strictly decrease with distance from the source, i.e., the maximal value occurs only at *x* = 0, whereas above the transition the marginal returns take on a constant value over the region 0 ≤ *x* ≤ *x*_0_ where enzymes are present and decreases only for *x* > *x*_0_. The level of this marginal returns plateau corresponds to the marginal cost *λ*_*e*_ of adding extra enzymes into the system (Fig. 2C, middle). This marginal cost decreases with increasing availability of enzymes (increasing *E_T_*). Fig. 2C shows how the shape of the marginal returns landscape changes when the enzyme profile deviates from the optimum. If it deviates by having a larger proportion of enzymes located at the source (Fig. 2C, top), the marginal returns landscape becomes non-monotonic with a peak at a position *x* > 0. The elevated enzyme density in this region depletes the substrate locally, reducing the turnover by each enzyme and diminishing the returns from adding additional enzymes. The peak in the marginal returns landscape implies that a higher flux can be achieved by moving some enzymes away from the source. In contrast, if the enzyme profile is overextended (Fig. 2C, bottom), the marginal returns landscape has a maximum at the source, such that the flux can be increased by moving enzymes towards the source.

Note that for the 1D problem analyzed in Fig. 2, the single source at *x* = 0 is always the position with the highest substrate concentration, independently of the enzyme arrangement (see purple lines). Hence an individual enzyme would always optimize its own productivity at *x* = 0. However, for *αE_T_* > 1, the total productivity of the population of enzymes is optimized only when some enzymes are located at positions *x* > 0 with non-maximal substrate concentrations. The optimal profile is thus the result of a bet-hedging strategy: Some productivity of individual enzymes is sacrificed to optimize the reaction flux produced by the entire population of enzymes. How relevant is the transition for biological systems? Given that the ratio *k*_cat_/*K*_M_ varies by orders of magnitude, 10^2^-10^9^(*Ms*)^-1^ [45], the corresponding *α* = *Lk*_cat_/(*K*_M_*D*) for a system with *L* = 1*μm*, *D* = 100*μm*^2^/*s* is in the range from 10^-7^ to 1, such that the transition can occur for enzyme numbers 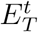 in the range 1-10^7^. For comparison, the range of protein copy numbers in a yeast cell is 1-10^6^ [46], suggesting that different enzymatic systems cover both sides of the transition.

### General condition for transitions in the optimal enzyme arrangement

Transitions in the optimal enzyme arrangement of the above type, from a regime in which enzymes are colocalized with the source of 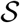 to a regime where enzymes are distributed within the system, have been observed in a range of reaction-diffusion models [18]. We therefore asked whether there was an underlying principle determining when such transitions occur, and whether it can be generalized, e.g. to systems with more complex geometries.

Fig. 2 suggests that the marginal returns landscape leads to a general condition for localization-delocalization transitions. For *E_T_* below the transition value, the landscape is strictly decreasing, with the position generating the highest returns coinciding with the source of substrate. As the transition is reached and passed, the landscape becomes flat as positions in the vicinity of the source begin to generate the same returns as the source position. This behavior generalizes to higher dimensional systems (see Appendix D): Enzymes should be placed only at the source on the surface and not in the interior, as long as

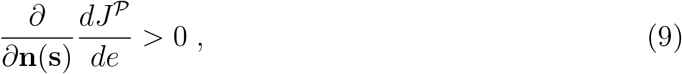

corresponding to a negative slope of the marginal returns landscape from the source at **s** into the interior of the system, as in Fig. 2B for *E_T_* < *α*^-1^. By contrast, if Eq. 9 evaluated at **s** becomes an equality, the positions adjacent to **s** generate the same returns as **s**, so that the optimal enzyme profile features enzymes both at the boundary **s** and in the interior of the system adjacent to **s**.

The abstract mathematical condition Eq. 9 can be turned into an experimentally meaningful condition (see Appendix D) by expressing it as a comparison between the local net diffusive flux *j^D^*(**s**) of 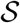 away from the surface, and the local flux 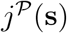 of product formation due to surface enzymes,

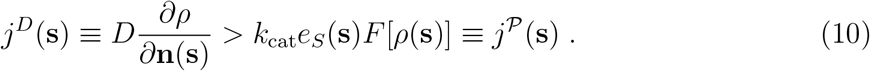

If this inequality holds, no enzymes should be placed in the interior of the system in the vicinity of **s**. For low amounts of enzymes, the optimal strategy is to place enzymes at position **s** to counter the diffusive flux *j^D^*(**s**). This is optimal up to the point at which the diffusive flux *j^D^*(**s**) equals the reaction flux 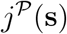, which defines the transition point of the localization-delocalization transition.

Further combining Eq. 10 with the boundary condition Eq. 2 yields 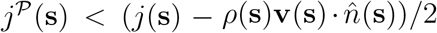, which simplifies in the case of no advection (**v** = 0) to 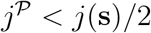. Thus, in systems with purely diffusive 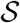 transport, a localization-delocalization transition occurs when the reaction flux equals exactly half the influx of 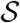. Note that Eq. 10 is general, independent of **v** and *σ* and other details of the system in question. It is required only that the boundary with influx follows Eq. 2. Other aspects of the model will affect *ρ*(**r**), but not the form of Eq. 10, showing that the diffusive motion of 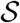 and the strength of the reaction are the crucial determinants of the transition point. To illustrate this generality, we compared the fluxes *j^D^*(**s**) and 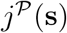 as *E_T_* was varied, in systems with different spatial dimensions as well as systems with and without decay and advection of 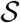. As shown in Fig. 3, in all cases both Eq. 10 below the transition value of *E_T_* and equality above the transition were satisfied.

**Figure 3.**
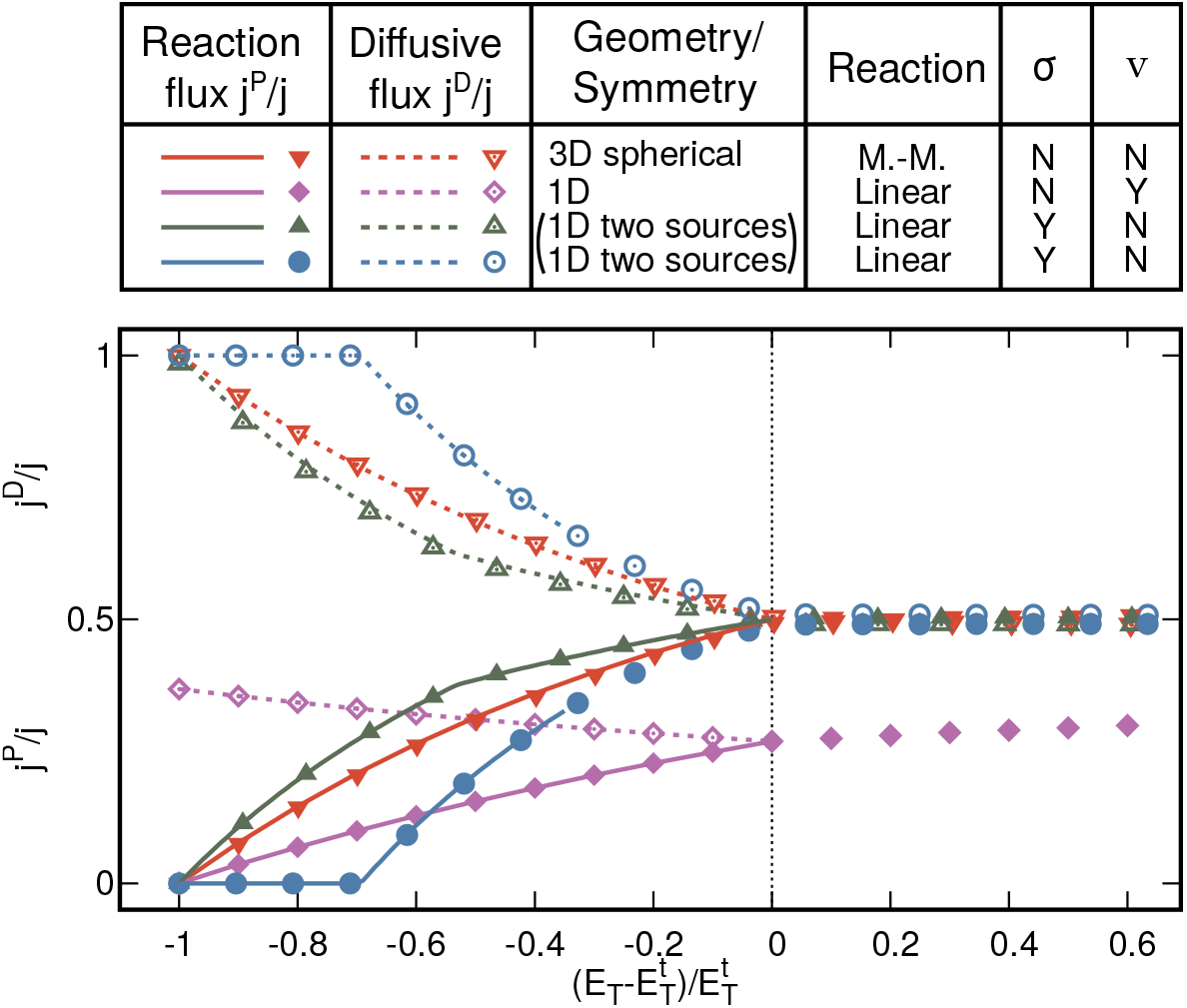
The local reaction flux, 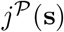, and net diffusive flux, *j^D^*(**s**), as a fraction of 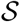 influx, *j*(**s**), are plotted for different models against the amount of enzymes in the system, *E_T_*, rescaled by the transition amount, 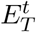. In all cases, 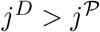 until they coincide at the transition point, as predicted by Eq. 10. When **v** = 0, 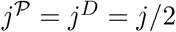 at and above the transition. Lines show analytical solutions of the constrained optimization. Points show results of numerical optimizations computed either by solving numerically an analytically-derived optimization condition (Appendix E), or using the construction algorithm. For the two-source model, the two curves show the two unequal sources, each rescaled with the corresponding 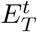 value; see Appendix F and Fig. F.1 for a detailed analysis of this model.

The inequality condition Eq. 10 is a strictly local condition. In systems with multiple or spatially varying sources of substrate there will in general not be a single global transition but rather a sequence of local transitions at different positions **s** at different *E_T_* levels. We therefore investigated a system with two unequal sources of 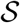. In such a system the optimal enzyme profile typically undergoes three transitions (see Fig. F.1). For small *E_T_*, enzymes accumulate where the influx of 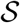 is the highest. As *E_T_* is increased, the marginal returns for placing enzymes in the vicinity of the two sources of 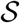 become more similar, until at a threshold value of *E_T_* they become equal. Above this value of *E_T_*, enzymes accumulate at both sources in a configuration with two unequal clusters. Increasing *E_T_* further, a second transition is reached where the optimal enzyme profile begins to extend from the stronger source into the system. Finally, at yet larger *E_T_*, the optimal enzyme profile begins to extend also from the weaker source. For the latter two transitions, at which the optimal enzyme arrangement changes from membrane-localized to extended, we confirmed that the inequality Eq. 10 held below the transition point and equality above (Fig. 3 green and blue).

### Geometrical interpretation and analogy to portfolio selection

Our optimal enzyme allocation problem has a geometrical interpretation in the space of (discretized) enzyme configurations **e**. A component *e_i_* of the vector **e** corresponds to the amount of enzymes allocated to position *i*, and the dimension of **e** to the number of possible enzyme positions. Hence, the subspace with constant total enzyme amount *E_T_* is the hyperplane intersecting each axis at *e_i_* = *E_T_*. Each point in the enzyme configuration space has an associated flux 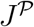, and the allocation problem amounts to finding the point of maximal flux within the hyperplane.

Fig. 4 illustrates the geometrical problem for the simplest case with only two possible enzyme positions (here, we assume a substrate source at position one, and loss of substrate via one of the loss mechanisms of Fig. 1B). In Fig. 4, *E_T_* hyperplanes are indicated by dashed black lines, while the dashed green curves are lines of constant flux 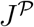. Since the marginal returns vector 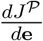 is the flux gradient, the lines of constant flux must be locally perpendicular to it. Adding extra enzymes at any location with nonzero substrate concentration will increase the total reaction flux, implying that the direction of the marginal returns vector always lies in the first quadrant (all components positive). Furthermore, the direction must turn clockwise as we move along a line of constant flux in the direction of increasing e_2_, since the second component of the marginal returns vector must decrease as we allocate a larger fraction of enzymes to position two, while the first component must increase. Hence, the lines of constant flux are convex.

**Figure 4.**
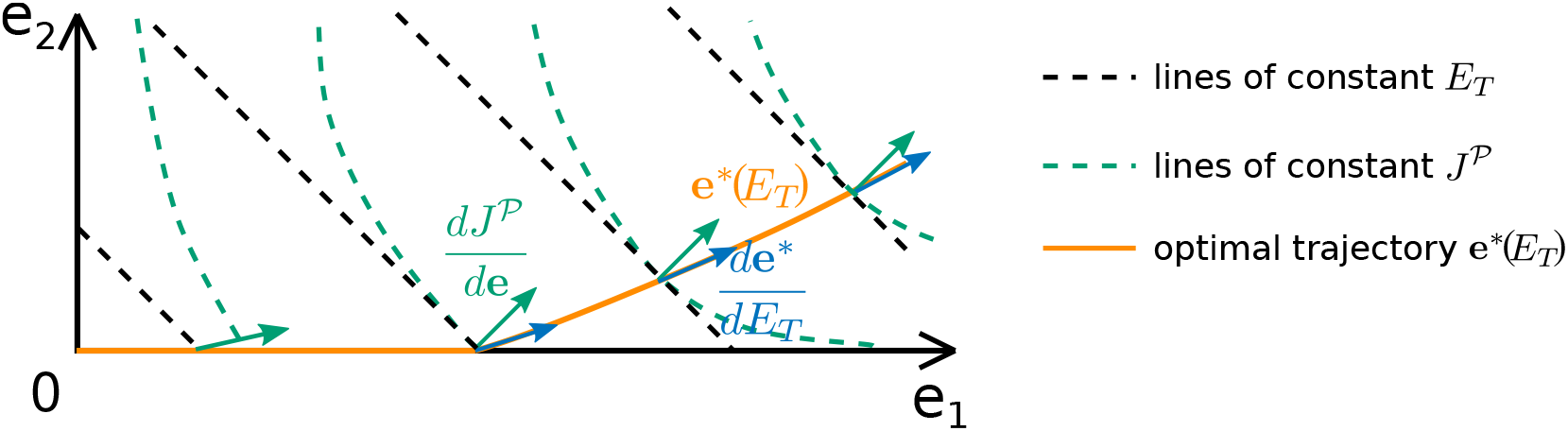
Geometrical analysis of the optimal enzyme configuration in a system with two sites. Dashed black lines show lines of constant *E_T_* = *e*_1_ + *e*_2_, while dashed green lines show lines of constant reaction flux 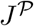. The optimal configuration 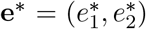 for each value of *E_T_* forms an optimal trajectory **e\***(*E_T_*) (orange). Green arrows show the marginal returns vector 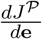. The tangent vector 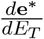 (blue arrows) shows how added enzymes should be optimally partitioned betweeen the two sites.

Geometrically, the solution of the optimal allocation problem corresponds to the touching point **e**\* between a given *E_T_* line and the 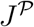 line with the largest 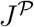 value that still touches the *E_T_* line. At small *E_T_* values, the 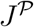 line is steeper than the *E_T_* line, such that **e\***(*E_T_*) lies on the **e**_1_ axis (orange line in Fig. 4). However, at a certain threshold value of *E_T_*, the *E_T_* line becomes tangential to the 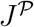 line. At this point, the marginal returns from position 1 and 2 become equal, i.e., the marginal returns vector points in the (1,1) direction. When *E_T_* is further increased, the optimal path **e\***(*E_T_*) departs from the *e*_1_ axis and follows the tangent point between the *E_T_* and 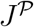 lines, maintaining equal marginal returns. Note that, due to the convexity of the lines of constant 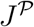, this tangent point is the global optimum at a given *E_T_* value.

The above geometrical interpretation is somewhat analogous to Markowitz’ geometrical analysis of investment portfolio optimization [11]. Markowitz considered the problem of minimizing the variance of a portfolio of investments at a constant level of returns (returns-variance analysis). The space of portfolios, which is determined by the relative amounts invested in different securities, is analogous to the space of enzyme configurations **e**. Markovitz noted that optimal portfolios are tangent points between lines of constant expected returns and lines of constant variance. One key difference to our case is that the returns are independent of the portfolio choice in Markovitz’ case, leading to simple straight lines of constant returns in the space of portfolios. Another key difference is that the variance of the returns has played no role in our analysis so far, since we considered enzymes in a steady-state environment. If, instead, we would consider a fluctuating environment in which, for instance, the substrate source is only available for a finite time *T*, then the variation of the flux 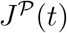 would become relevant. The total amount of product formed during this time, 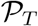, would be

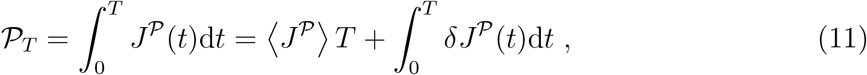

where 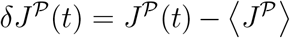 is the time-dependent fluctuation of the flux around its mean value 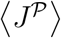, caused by the stochasticity of the enzymatic reaction and substrate diffusion. The integral over 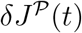 scales as 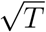, due to the central limit theorem, such that its contribution to 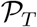 becomes negligible in the long time limit. However, for times *T* comparable to the intrinsic time-scales of the system (associated with the processes of diffusion, reaction, and decay), the contribution of the fluctuations could become sizeable. In such scenarios, a generalization of Markovitz’ returns-variance analysis would be more appropriate than an optimization of 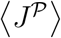.

### Additive construction of optimal enzyme arrangements

The above geometrical analysis leads us to an additive construction scheme, which generates the trajectory of optimal enzyme configurations as enzymes are added to the system. Generalizing from the example of Fig. 4, we consider a system in which space is discretized into *N* sites, denoting by *e_i_* = *e*(**r**) the density of enzymes at position **r** and by *ρ_i_* = *ρ*(**r**) the corresponding substrate density. For a given total enzyme level *E_T_*, there will be an optimal enzyme configuration **e**\* = 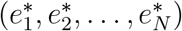) that maximizes 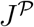 on the (*N* – 1)-dimensional hyperplane defined by 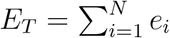. As in Fig. 4, the optimal configurations at different *E_T_* values trace out a trajectory **e\***(*E_T_*) in **e** space. We seek a procedure to construct **e\***(*E_T_*). We begin with no enzymes and iteratively add enzymes until the target *E_T_* value is reached. According to the principle of homogeneous marginal returns, the largest components of 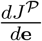 define the sites at which enzymes should be added in the next step. Addition may initially be restricted to a single site, with further “construction sites” being introduced gradually as different *E_T_* thresholds are crossed.

Since enzymes should be added at all positions where the marginal returns take on their maximal value, we must determine how to partition new enzymes between these positions. This partitioning corresponds to the tangent vector 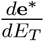 of the optimal trajectory, which can be obtained from the variation of the marginal returns as enzymes are added (see Appendix G1),

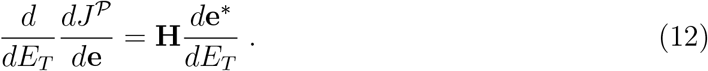

Here, derivatives 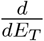 are taken tangentially to the line of optimal enzyme configurations and **H** is the *N* × *N* Hessian matrix of 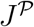 in the space of enzyme densities, 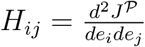. In the subspace of only those sites where enzymes should be added (sites at which 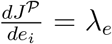), the left hand side of Eq. 12 becomes

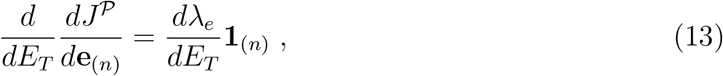

where the subscript (*n*) indicates that we are considering only the *n*-dimensional subspace of sites with equal returns, yielding

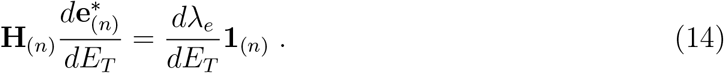

This *n*-dimensional linear system can be solved for the tangent 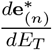 to the optimal trajectory [47]. Taken together, we obtain the following additive construction scheme for the optimal enzyme arrangement:

1. Begin with **e** = 0.
2. For each site **r**_*i*_, evaluate the marginal returns 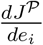 for adding enzymes at that position.
3. Identify the subset of positions at which the returns are equal (within numerical tolerance) to the maximal value.
4. For this subset of positions, evaluate the Hessian matrix **H**(_*n*_) of the flux with respect to the enzyme densities.
5. Solve **H**(_*n*_)*δ***e**\* = **1** for *δ***e**\*, and normalize such that the total change in the amount of enzymes 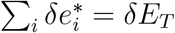.
6. Update the optimal enzyme arrangement, **e**\* → **e**\* + *δ***e\***.
7. Repeat from step 2 until the target total enzyme amount *E_T_* is reached.

The discretization of the system, which is required for the above construction algorithm, can be chosen using a criterion based on Eq. 10 (see Appendix G2).

Fig. 5 illustrates the algorithm for a two-dimensional system featuring multiple sources of substrate (dashed rectangles) with uniform influx on their surfaces and an absorbing outer boundary. Fig. 5A compares two optimal enzyme configurations with a different amount of enzymes. For low *E_T_* values, enzymes localize non-uniformly at the boundaries of the sources. For higher *E_T_* values, the optimal enzyme profile extends into the interior of the system. However, the highest densities of enzymes are found in the regions between the sources and the outer boundary of the system, where the gradient of substrate concentration is steepest. A significant fraction of enzymes are thus devoted to substrates that, by the nature of where they are produced, are likely to rapidly diffuse out of the system.

**Figure 5.**
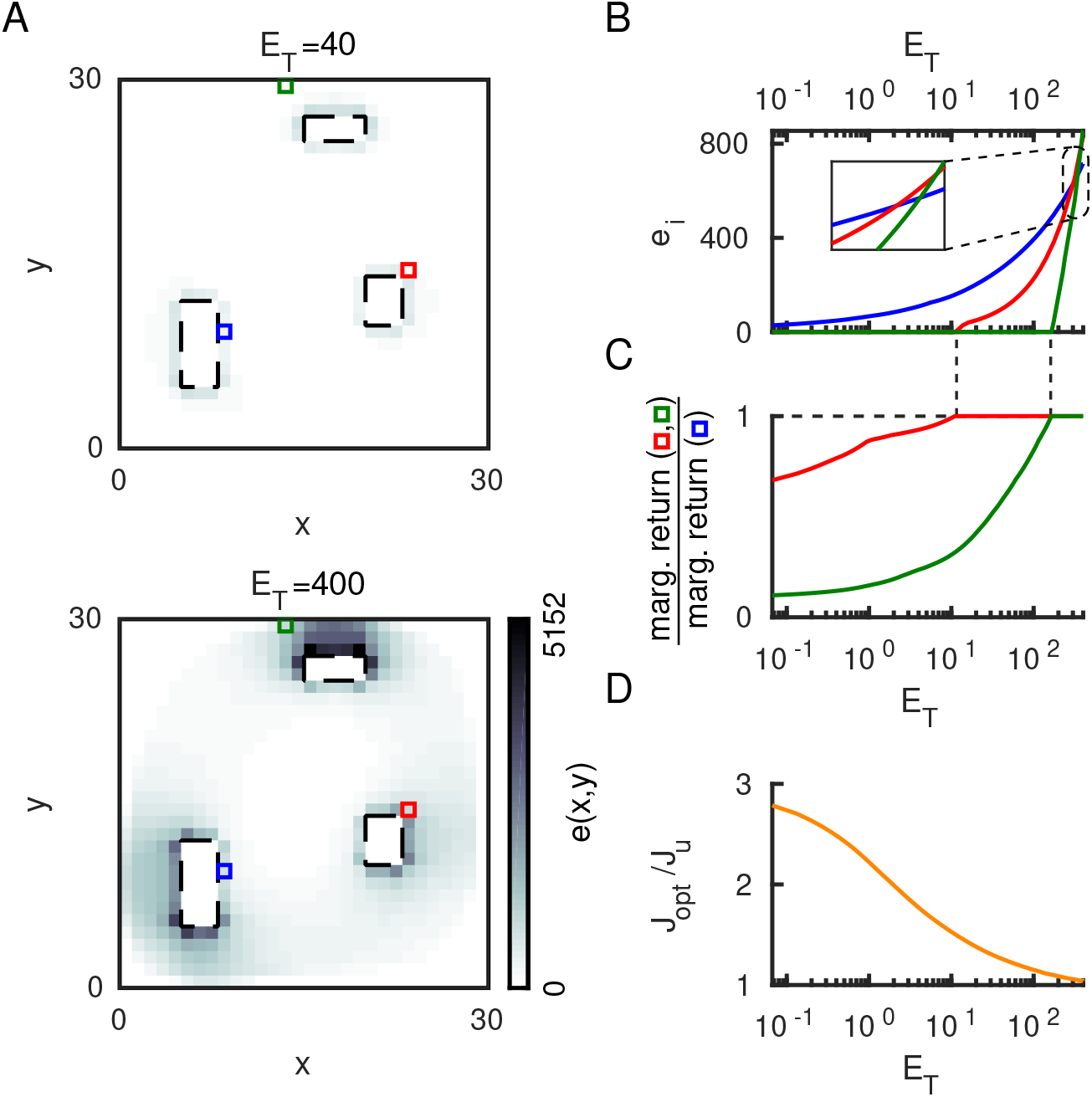
Optimal enzyme arrangements in a 2-dimensional system with multiple sources of substrate (dashed rectangles), calculated using the construction algorithm. (A) Snapshots of the optimal enzyme distribution at two *E_T_* levels. (B) Optimal enzyme densities and (C) ratios of marginal returns at the positions indicated by colored squares in (A). (D) The optimized reaction flux *J*_opt_ relative to the flux *J*_u_ produced by uniformly distributed enzymes, as a function of *E_T_*.

Remarkably, the sites with the highest enzyme densities at small *E_T_* do not necessarily retain the highest densities as *E_T_* is increased. Fig. 5B compares the enzyme densities as a function of *E_T_* at three positions (marked by colored squares in Fig. 5A). The blue location is the first at which enzymes are added. However, although enzymes are added at the green and red positions only later, the densities here become higher. Fig. 5C shows the respective ratios of the returns of the red and green position to the returns from the blue position. Enzymes are added at the green and red position only when the returns are equal and the ratios are one. At low enzyme amounts, the reaction flux of the optimal configuration is about 3-fold higher than for the uniform configuration, whereas this advantage gradually decreases for higher amounts of enzyme (Fig. 5D).

Compared to the stochastic search algorithms that have previously been used to find the optimal enzyme arrangements in systems with simple geometries [17, 18], the additive construction scheme derived here has several advantages. In terms of computational effort, it generally performs better than stochastic optimization (see Appendix G3), in particular for more complex systems. Moreover, while stochastic optimization proceeds by trial-and-error and can get stuck in local optima, the construction scheme follows a rational principle. By design, the construction scheme not only generates the optimal enzyme arrangement for the given total enzyme amount *E_T_*, but all optimal configurations for total enzyme amounts up to this level. As seen in the example of Fig. 5, this trajectory of optimal configurations leads to additional insight about the properties of the particular system at hand.

The construction scheme can also be adapted to other allocation problems in which investments and returns are coupled. As long as the resources to be invested follow a linear budget constraint, the adaptation amounts to a modification of the objective function. For instance, as mentioned above, one could consider an objective function penalizing high variance as in Markovitz’ problem [11]. One could also consider problems where the capital of the investor grows exponentially (iterative reinvestment) as in Kelly’s problem [10] or more complex variants [48]. In each case, one needs to compute the marginal returns and the Hessian of the new objective function with respect to the resources invested in the different assets, and determine the optimal allocation of resources as the total budget gradually increases via the construction scheme. In Appendix H, we present a generalization of Kelly’s problem to a case with coupled investments and returns for which the objective function becomes the long-term capital growth rate and the above additive construction scheme can be used.

## DISCUSSION

We presented a solution for the problem of optimally allocating enzymes in space, to maximize the reaction flux of an enzymatic reaction. Our solution encompasses a class of systems with potentially complex geometries, in which the substrate enters via internal or external boundaries, is transported via diffusion and possibly advection, and can be lost by leakage or competing reactions (Fig. 1). The solution is based on the concept of a ‘marginal returns landscape’ (Fig. 2), defined as the derivative 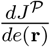 of the total reaction flux with respect to the local enzyme density (Eq. 7). The optimal arrangement of enzymes is such that the marginal returns are spatially homogeneous over all positions with enzymes. This criterion of homogeneous marginal returns (HMR) can more generally be applied to betting games where each bet globally feeds back onto all returns (Appendix H). In the enzyme allocation problem, the feedback is generated by enzymes locally depleting their substrate, which affects the global substrate profile.

The HMR criterion leads to a general local condition for the occurrence of a localization-delocalization transition in the optimal enzyme arrangement (Eq. 10), which compares the local diffusive flux *j^D^*(**s**) to the local reaction flux 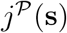 at a source position **s**: As long as 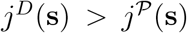, localization of enzymes at the source is optimal, while the two fluxes are equal at the transition point and above (Fig. 3). The simplicity of this flux condition makes it practical for application to experimental systems. For instance, for an *in vivo* system in which enzyme localization is observed and the enzyme abundance is quantified, the condition can be used to estimate whether the observed enzyme localization indeed maximizes the reaction flux, or whether it is more likely to play a regulatory role [7, 8]. In this context, it is beneficial that the flux condition is not affected by any advective transport of the substrate, which may be present in the system, e.g. due to cytoplasmic streaming or transport by molecular motors. This independence is due to the deterministic nature of advective transport - the HMR criterion is fundamentally a bet-hedging strategy to deal with the probabilistic nature of diffusive transport.

Just as the Kelly criterion [10] and returns-variance analysis [11] provide rules for optimally placing bets or optimally constructing portfolios, the HMR criterion allowed us to derive an algorithm for constructing optimal enzyme arrangements (Figs. 4 and 5). This algorithm is obtained by analyzing how the marginal returns change at each position as the amount of enzymes in the system is increased (Eq. 14). It constitutes an exact deterministic construction scheme, which is applicable to systems with arbitrary geometries and multiple sources. Here, we presented the simplest general form of the algorithm. Various modifications can be made to improve its efficiency for particular systems. Using the system geometry as a guide, one can limit the sites at which the marginal returns need to be evaluated in each update step. For example, at low *E_T_* values the positions with the highest returns coincide with sources of 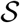. Generally it suffices to compute the marginal returns only at source positions, sites where enzymes are already present, and neighboring sites. One could further speed up the algorithm by enabling larger increments *δE_T_* with higher-order update schemes for **e**\*.

We explicitly considered absorbing and reflective boundaries, also allowing for non-uniform and discontinuous influx profiles. However our results can be extended to systems with partially permeable boundaries, corresponding to *k*(**s**) = 1 and *h*(**s**) = *p*(**s**) > 0 in Eq. 3, where *p*(**s**) is the permeability. For such systems, the HMR criterion still holds. Similarly, the condition of Eq. 9 still determines transitions in the optimal enzyme profile. However, expressing this condition in term of substrate fluxes will lead to a more complicated expression than Eq. 10. A permeable boundary typically increases the transition threshold 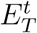. For example, in the one-dimensional system of Fig. 2, a non-zero permeability at the origin shifts the threshold value of *E_T_* from *α*^-1^ to *α* ^-1^(1 + *p*(0)*L/D*).

Our continuous reaction-diffusion model neglects the finite size of enzyme molecules that limits the attainable enzyme density. Imposing a maximal density condition *e*(**r**) ≤ *e*_max_ at each point [16] could be incorporated into our analytical framework as an additional constraint in the Lagrangian. This would introduce an effective position-dependent cost in Eq. 7, resulting in a marginal returns landscape that is a function of position. The construction algorithm could still be employed with the additional condition that enzymes cannot be added at positions where *e_i_* = *e*_max_, such that Eq. 14 would be restricted to positions with sub-maximal enzyme densities.

The construction algorithm for optimal enzyme arrangements appears suited for applications in synthetic biology [19]. The attachment of enzymes to membranes can be controlled by fusing enzymes to transmembrane proteins [49, 50] or by using protein scaffolds [51, 52]. The arrangement of enzymes in the interior of membrane-enclosed systems can be partially controlled with RNA assemblies [53], synthetic protein scaffolds [20], or fusion proteins [16]. Enzyme positioning on surfaces can be controlled with nanometer precision via single-molecule cut-and-paste surface assembly [54, 55], and in 3D by arranging enzymes on DNA scaffolds [56–59]. Combining this arsenal of techniques with the rational construction scheme, one could optimize synthetic bioreactors to produce drug molecules [20–22] or biofuels [23, 24].

## MATERIALS AND METHODS

### Computation of 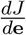 H for linear reactions

For a discrete reaction-diffusion system with linear reactions, following the dynamics of Eq. 1, the steady-state equation can be written in matrix form as

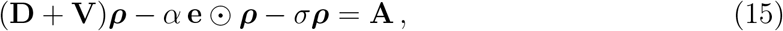

where **D** and **V** are the diffusion and advection operator on the lattice respectively, ***ρ*** the substrate density and **A** the substrate influx vector, while Θ denotes the element-wise (Hadamard) product. The reaction flux 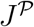 is given by the scalar product 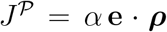. Thus **e** enters 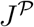 both directly and through the solution ***ρ*** of Eq. 15. These multiple dependencies of 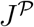 on **e** are summarized by the dependency graph in Fig. 6, where the matrix **M** = **D** + **V** – *α* diag(**e**) – *σ* diag(**1**) is the operator that applied to ***ρ*** gives the substrate source vector **M_ρ_** = **A**, diag(**e**) is the diagonal matrix with diagonal elements given by **e** and diag(**1**) is the identity matrix.

**Figure 6.**
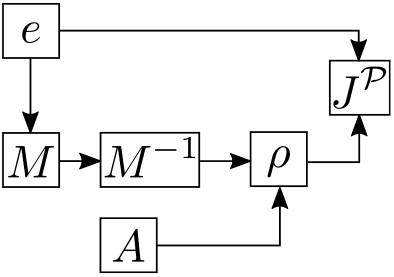
Dependency graph for a discrete reaction-diffusion system with linear reaction kinetics.

The marginal returns vector 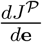 can be evaluated by back-propagating the derivative of the reaction flux with respect to the enzyme vector through the dependency graph. We find that

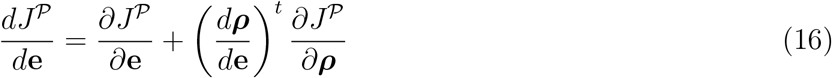

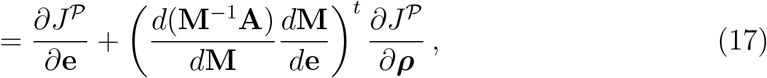

where 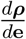 is a matrix with (*i, j*)^th^ element given by 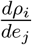, and ^*t*^ denotes the transpose. Each of the derivatives appearing in Eq. 17 has a simple form, which we substitute to obtain the marginal returns vector,

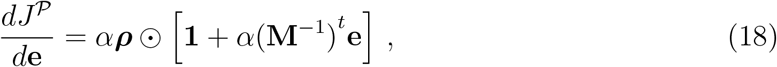

where **1** is a vector of ones.

By taking a second derivative with respect to **e**, and again by back-propagating the derivatives, we can determine the Hessian,

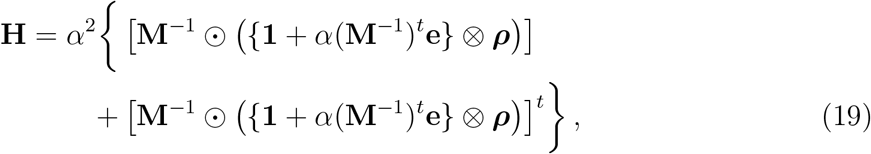

where ⊗ indicates the outer product.

### Non-linear reactions

For non-linear reaction kinetics the reaction-diffusion equation becomes

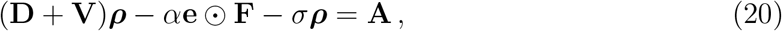

where **F** is the vector with *F_i_* = *F*[*ρ_i_*]. Since *F*[*ρ*] is non-linear, it is no longer possible to write the reaction-diffusion equation as a linear system that can be solved for ***ρ***. As a result the dependency graph includes a loop, making it inconvenient to back-propagate derivatives. Instead, taking derivatives of the reaction-diffusion equation with respect to **e**, we can directly solve for

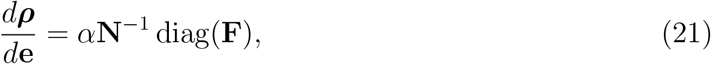

where **N** = **D** + **V** – *α* diag(**e** ⊙ **F**′) — *σ* diag(**1**) and 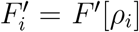. We then take derivatives of 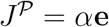. **F** with respect to **e**, to obtain the marginal returns vector,

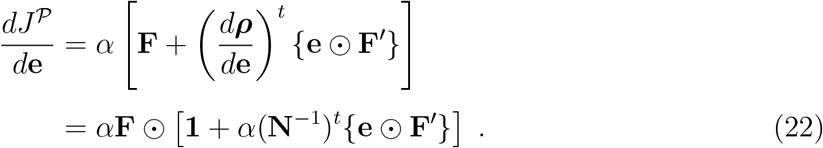

By taking a further derivative of the returns vector with respect to **e** we find the components of the Hessian,

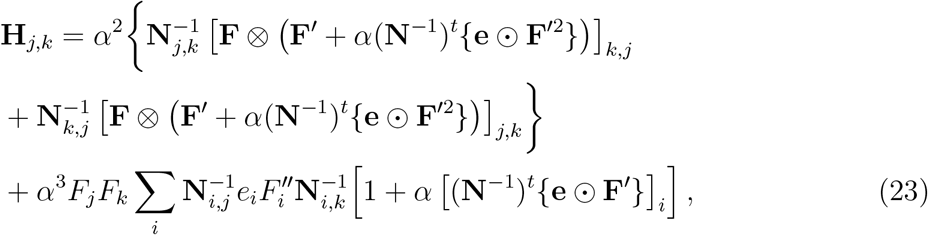

which simplifies to Eq. 19 for linear reactions with *F″* = 0.

## ACKNOWLEDGMENTS

This work was supported by the German Research Foundation via SFB1032 “Nanoagents for Spatiotemporal Control of Molecular and Cellular Reactions” and the excellence cluster ORIGINS. G.G. was supported by a DFG Fellowship through the Graduate School of Quantitative Biosciences Munich (QBM).

## AUTHOR CONTRIBUTIONS

All authors designed research and developed the theoretical framework. G.G. and F.T. developed computational methods and analyzed data. G.G., F.T., and U.G. wrote the paper.

## Appendix A: General Lagrangian with multiple subspaces

As noted in the main text, our functional Lagrangian formalism assumes smooth, positive enzyme distributions within different subdomains. At the interfaces of such domains, the optimal enzyme distributions may feature discontinuities. For example, there may be a finite enzyme density in one region of the system, and no enzymes (zero density) in another region. These subdomains should be picked in the proximities of the substrate sources. For example if we have patches of sources on the surface domains, for each source patch we would consider a surface subdomain around the patch and a correspondent volume subdomain in proximity of the source. These are the regions where it is optimal to distribute the enzymes as diffusion does not allow for peak of substrate concentration away from the sources. Within each such subdomain *i*, the substrate and enzyme densities *ρ*^(*i*)^(**r**) and *e*^(*i*)^ (**r**) are smooth and their local gradients are well defined. However, at the interfaces between such subdomains, *e*(**r**) need not be continuous (although *ρ*(**r**) still must be). The positions of the interfaces, and therefore the extents of the different domains, should be optimized over and as we have seen in the main text they depend on the *E_T_* considered.

In this formulation, the total reaction flux can be written as a sum of contributions from each surface and boundary domain, and the corresponding reaction-diffusion or boundary condition acts as a constraint that must be satisfied within each domain. In addition, we introduce explicit matching conditions on *ρ*(**r**) at the interfaces *I*. The resulting Lagrangian has the following form:

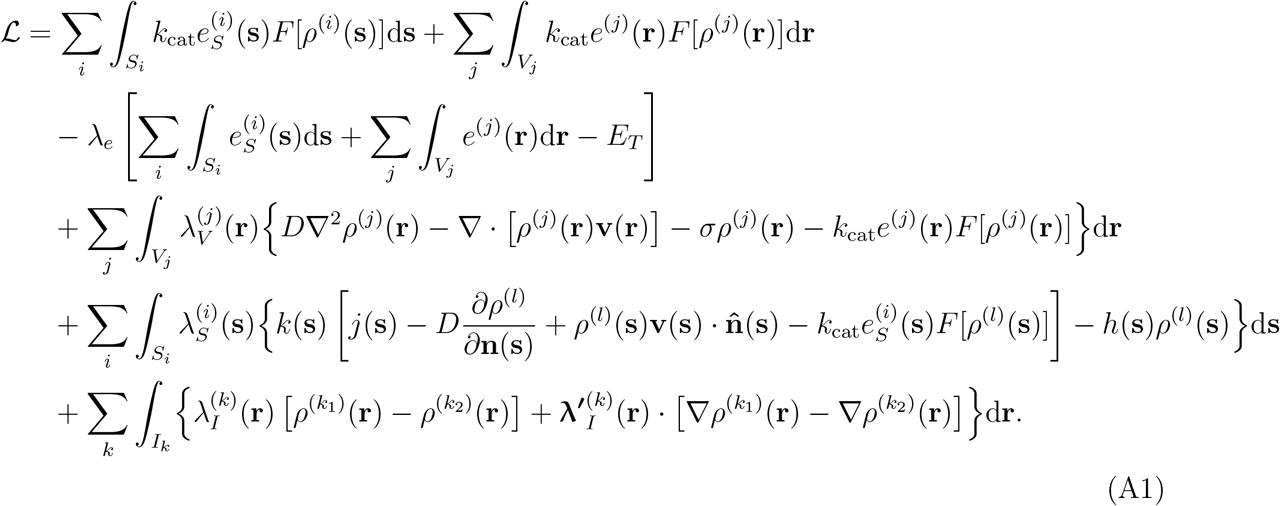

where *l* represents the index of the volume domain adjacent to the boundary at **s**, and *k*_1_, *k*_2_ represent the two domains on either side of the interface *I_k_*. Here *λ_I_*(**r**) is a Lagrange multiplier that enforces smoothness of the density profile at each point **r** on an interface between two domains, and 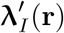 is a vector of Lagrangian multipliers where each component enforces smoothness of *ρ*(**r**) along one of the system dimensions.

It is convenient to consider first some manipulations of the general Lagrangian, Eq. A1. Using the vector identity ∇ · (*f***a**) = ∇*f* · **a** + *f*∇ · **a** we can rearrange the diffusive and advective transport terms to read

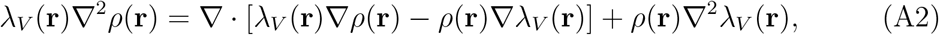

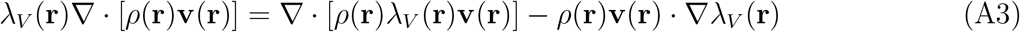

Applying Gauss theorem (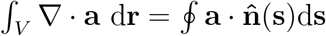), Eq. A1 becomes

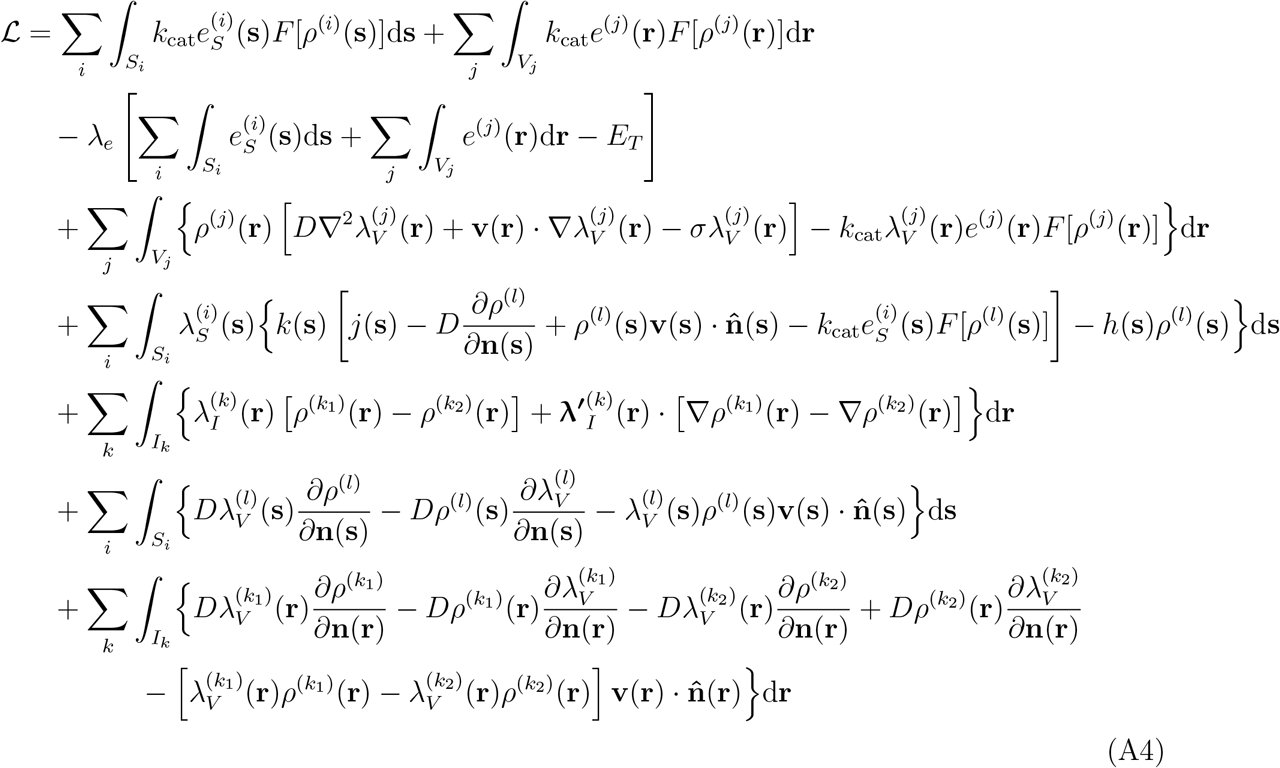

Here the last three lines contain the contributions from the boundary integral.

## Appendix B: Optimal enzyme arrangement in a one-dimensional system

In this section we go through in detail the full analytic optimization of the simplest exactly-solvable model. We consider a one-dimensional system with a source of substrate at one boundary at *x* = 0, (*j*(0) = *j*_0_, *k*(0) = 1, *h*(0) = 0) and an absorbing boundary at *x* = *L* (*j*(*L*) = *k*(*L*) = 0, *h*(*L*) = 1). Further we assume a linear reaction functional, no drift and no instability of 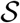, i.e. *F*[*ρ*] = *ρ*/*K*_M_ and *v* = *σ* = 0.

In our formalism we divide the system into two boundary domains (*S*_1_: *x* = 0 and *S*_2_: *x* = *L*) and two “volume” domains (*V*_1_: 0 ≤ *x* ≤ *x*_0_ and *V*_2_: *x*_0_ ≤ *x* ≤ *L*, with 0 ≤ *x*_0_ ≤ *L* a parameter to be optimized). For simplicity of presentation we will assume that enzymes can be present at *x* = 0 or in the domain *V*_1_, while the boundary at *x* = *L* and the domain *V*_2_ are free of enzymes. Whilst one can include an arbitrary number of additional domains within the system, we found that such domains turned out to be degenerate with the two specified above. Therefore, we will not include these possibilities here. The effective enzyme profile is therefore of the form *e_S_δ*(*x*) + *e*(*x*)Θ(*x*_0_ – *x*), where *δ*(*x*) is the Dirac delta function and Θ(*x*) is the Heaviside function.

The Lagrangian for this specific model is

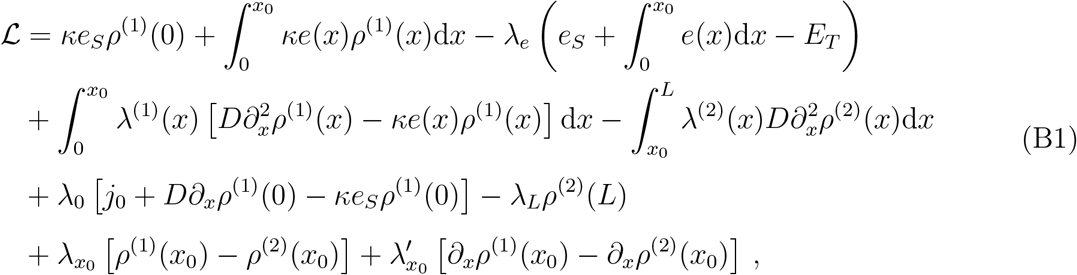

where we have introduced the catalytic efficiency *κ* = *k*_cat_/*K*_M_. Here *λ_e_* is the Lagrange multiplier corresponding to the constraint of having a fixed amount of enzymes *E_T_*; λ^1)^(*x*) and λ^(2)^(*x*) those corresponding the constrained dynamics of the substrate in the two domains; λ_0_ and λ_*L*_ to the boundary conditions; and λ_*x*0_ and 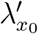 to the continuity and smoothness that must be imposed on *ρ*(*x*) at the interface *x*_0_.

To calculate the optimal enzyme arrangement 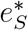, *e*\*(*x*) we must compute the various functional derivatives of 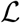. Let us first consider the derivatives with respect to *e*(*x*) and *ρ*^(1)^(*x*). After integrating by parts Eq. B1 (corresponding to the form Eq. A4 of the Lagrangian), we find

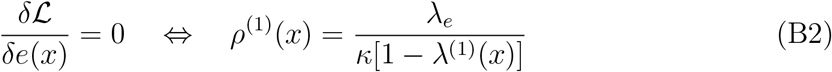

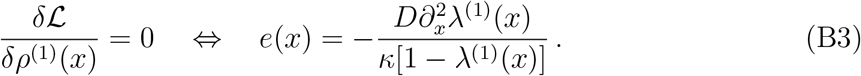

We can compute the derivatives *∂_x_ρ*^(1)^(*x*) and 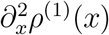 starting from the expressions above and we can substitute the results into the reaction-diffusion equation (given also by 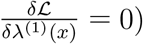 to obtain an ordinary differential equation for λ^(1)^(*x*):

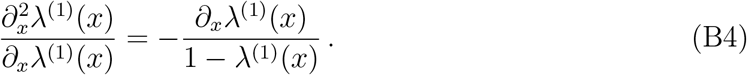

The solution to Eq. B4 can be found by direct integration to be

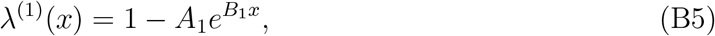

where *A*_1_ and *B*_1_ are constants of integration still to be determined. Using Eqs. B2-B3 we obtain

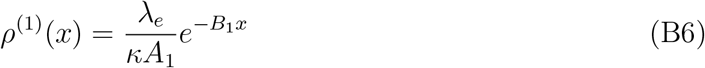

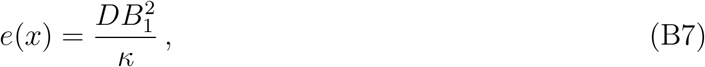

Equation B7 immediately shows that the enzyme density in the region 0 ≤ *x* < *x*_0_ is constant, in agreement with previous numerical studies [17, 18].

Evaluating the functional derivative of 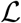 with respect to λ^(2)^, we recover the diffusion equation 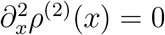 in the domain *V*_2_, from which

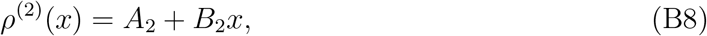

where *A*_2_ and *B*_2_ are again constants of integration.

The constants *A*_1,2_ and *B*_1,2_ can be evaluated using the boundary conditions at *x* = 0 and *x* = *L*, and the continuity and smoothness conditions at *x* = *x*_0_. Additionally, the dependence on *e_S_* can be eliminated using 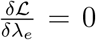, which is simply the constraint on the total enzyme number. This leaves us with an expression for the Lagrangian that, since all constraints have been satisfied equals the constrained reaction flux 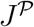, in terms of the single optimization variable *x*_0_,

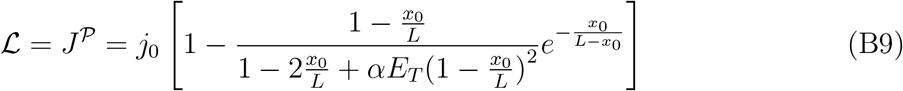

where we define *α* = *κL/D*. Finally, we maximize over *x*_0_ to find

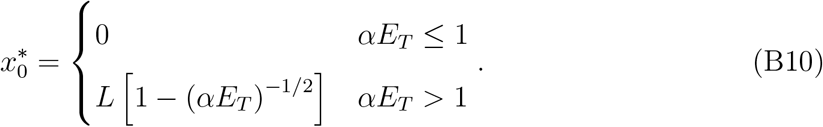

By substitution we can then obtain expressions for the enzyme and substrate densities

**Table B.1.** Optimal enzyme distribution and corresponding substrate profile and flux for a one-dimensional system

| Regime | $e_S^*$ | $e^*(x)$ | $\rho^{(1)}(x)$ | $\rho^{(2)}(x)$ | $J^{\mathcal{P}*}$ |
| --- | --- | --- | --- | --- | --- |
| $\alpha E_T \leq 1$ | $E_T$ | 0 | N/A | $\frac{j_0(L-x)}{D(1+\alpha E_T)}$ | $\frac{j_0}{1+(\alpha E_T)^{-1}}$ |
| $\alpha E_T > 1$ | $\sqrt{\frac{E_T}{\alpha}}$ | $\frac{E_T}{L}$ | $\frac{j_0}{2} \frac{L}{D\sqrt{\alpha E_T}} \exp\left[-\sqrt{\alpha E_T} \frac{x}{L}\right]$ | $\frac{j_0}{2} \frac{L-x}{D\sqrt{\alpha E_T}} \exp\left[-\sqrt{\alpha E_T} \frac{x_0}{L}\right]$ | $j_0 \left(1 - \frac{1}{2} e^{1-\sqrt{\alpha E_T}}\right)$ |

## Appendix C: Marginal returns landscape

Given a particular enzyme arrangement, the marginal returns as a function of position can be calculated directly by considering the addition of an amount *δe* of enzymes at position *x*′, and solving the new modified reaction-diffusion system. Here we present an example of such a calculation. We consider the same one-dimensional system analyzed above, and suppose that we are in the regime *αE_T_* > 1 and the enzyme profile is the optimal one of Table B.1. For other enzyme profiles, such as those shown in Fig. 2C of the main text, the calculation proceeds in the same way.

We now have to consider two cases: 0 ≤ *x*′ ≤ *x*_0_ and *x*_0_ < *x*′ ≤ *L*.

If 0 ≤ *x*′ < *x*_0_, we solve the reaction-diffusion equation in the three domains: 0 ≤ *x* < *x*′, *x*′ < *x* ≤ *x*_0_, and *x*_0_ ≤ *x* ≤ *L*. The resulting density profiles are

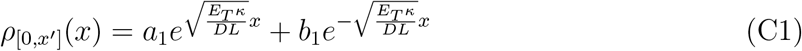

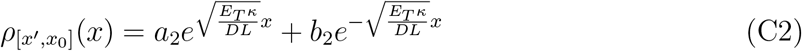

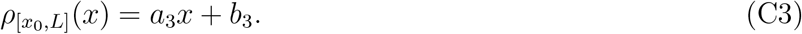

To fix the constants of integration *a_i_, b_i_*, we make use of the boundary conditions at *x* = 0 and *x* = *L* as well as matching conditions on *ρ* at *x*′ and *x*_0_. Specifically, we require continuity of *ρ* at *x*′ and *x*_0_, and smoothness at *x*_0_,

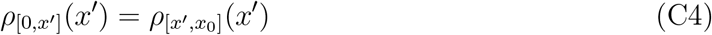

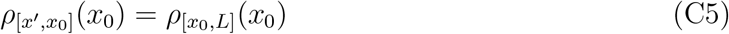

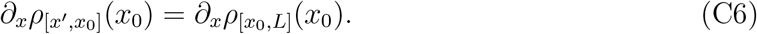

Additionally, conservation of mass at *x* = *x*′ requires that

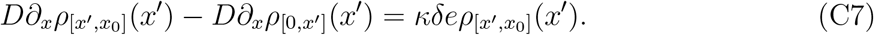

The total flux 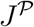 is then given by

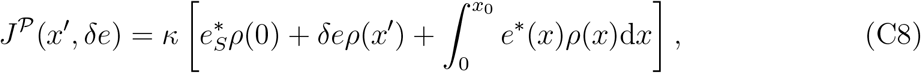

into which we substitute the final density profile *ρ*(*x*) as well as 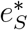 and *e*\*(*x*) from Table B.1.

Finally, the marginal returns at position *x*′ can be calculated as

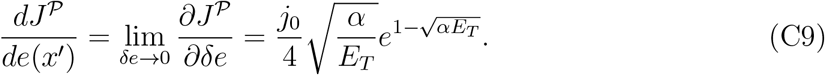

Therefore, for 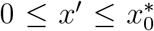 the marginal returns do not depend on *x*′, and all positions at which we have enzymes give the same returns. It is straightforward to verify that this agrees with the value of λ_*e*_ derived from the optimal reaction flux that appears in Table B.1,

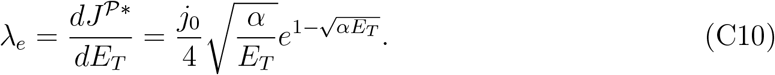

In the case *x*_0_ ≤ *x*′ < *L*, the same procedure can be applied, except with the three domains 0 ≤ *x* ≤ *x*_0_, *x*_0_ ≤ *x* ≤ *x*′, and *x*′ ≤ *x* ≤ *L*. We find that the marginal returns is given by

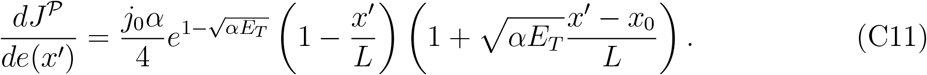

As expected, this approaches Eq. C9 as 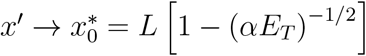.

## Appendix D: Derivation of the transition condition

In this section we derive the inequality condition, Eq. 10 of the main text, that determines whether or not enzymes should be placed in the volume adjacent to a boundary position **s**. We begin by examining the functional derivative of 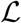 with respect to substrate profiles at the system boundary, *ρ*(**s**). In the vicinity of a reflecting boundary, (i.e. *h*(**s**) = 0, *k*(**s**) = 1), we obtain from Eq. A4

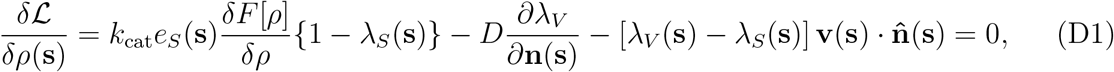

where equality with zero assumes that the enzyme distribution is the optimal one. We can also consider the functional derivative with respect to the gradient of *ρ*(**r**) at the boundaries of the system,

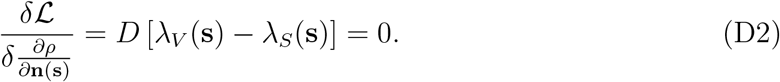

Equation D2 requires λ_*V*_(**s**) = λ_*S*_(**s**), which simplifies Eq. D1 to

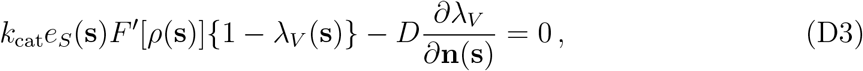

where 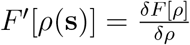.

We next examine variations of 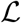 with respect to the enzyme densities,

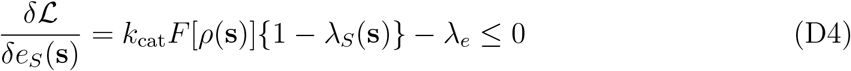

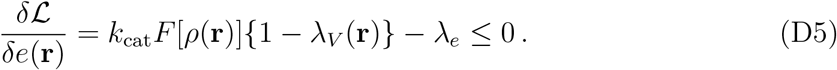

Both λ*V*(**r**) and *ρ*(**r**) must be continuous, since they must both satisfy (generally non-linear) diffusion equations. Therefore, it follows from Eq. D5 that 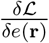 is itself continuous and that we can define the spatial gradient 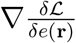. At all points where it is optimal to allocate a non-zero density of enzymes (both in the interior of the system or on the boundaries) we have that 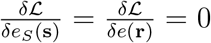, implying that 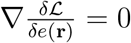. In regions where enzymes are not present because it is suboptimal to add them, we get that 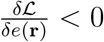, and the gradient 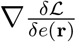 will in general be finite. Furthermore, since *ρ*(**r**) → *ρ*(**s**) and λ_*V*_(**r**) → λ_*V*_(**s**) = λ_*S*_(**s**) smoothly as **r** → **s**, 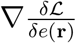 must remain well defined as we approach the system boundaries.

If it is optimal to have enzymes at the boundaries of the system but not locally in the interior, 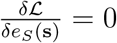 and 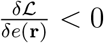 for a small displacement **r** = **s** + **ϵ**. In this case, evaluating the gradient of Eq. D5 and taking **r** → **s** we find

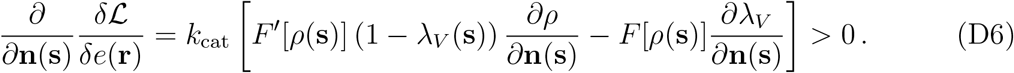

The inequality Eq. D6, 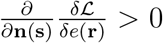, is equivalent to Eq. 9 of the main text, 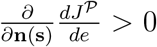, because 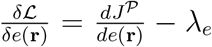 and λ_*e*_ is constant. Hence 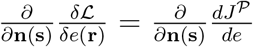. Substituting for 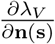 from Eq. D3, this inequality becomes

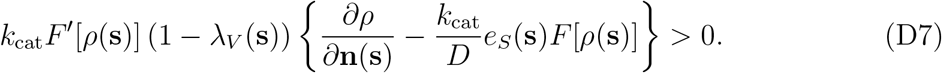

Since we assume that *F*[*ρ*] is monotonically increasing, *F*′[*ρ*] > 0. Furthermore, setting Eq. D4 equal to zero implies that 1 – λ_*V*_(**s**) > 0, since λ_*e*_ > 0. Therefore, Eq. D7 reduces to

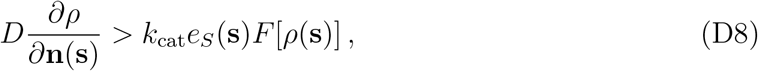

which corresponds to Eq. 10 of the main text.

## Appendix E: Reduction of numerical complexity in a one-dimensional system with drift

In this section we demonstrate how, even in cases for which the full analytical solution is not tractable, it is nevertheless possible to use the Lagrangian formalism to express the optimization problem as a single equation, which can then be solved numerically.

We consider a one-dimensional system, similar to that considered in Section B but with the addition of a constant drift *v*. If *v* > 0 substrate molecules tend to flow away from the source at the origin, whereas for *v* < 0 they tend to flow towards it. The corresponding Lagrangian is

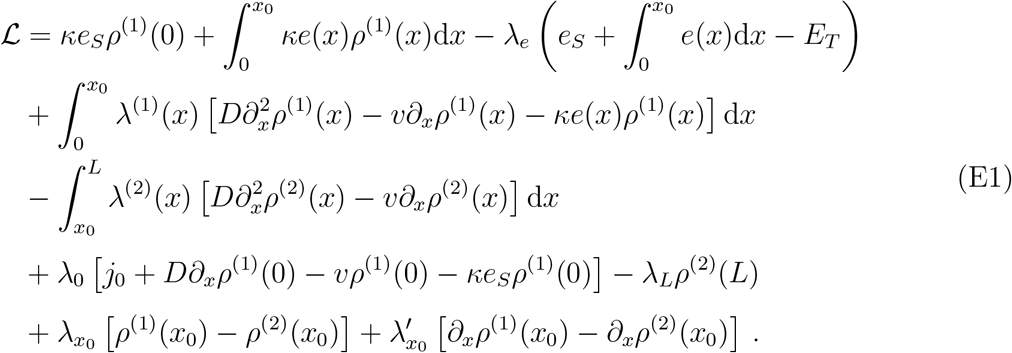

The derivation proceeds in the same way as in Section B. We again use the functional derivatives to find expressions for *e*(*x*) and *ρ*^(1)^(*x*) in terms of the Lagrange multiplier λ^(1)^ (*x*),

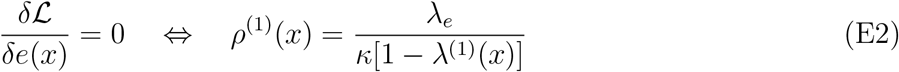

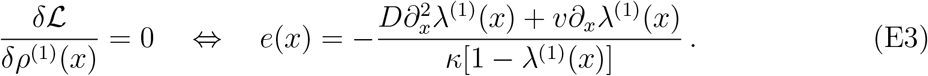

Surprisingly, when we combine these expressions with the reaction-diffusion equation with drift, we find that the resulting differential equation for λ^(1)^(*x*) is again exactly Eq. B4, which we can solve to find

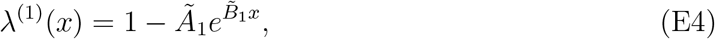

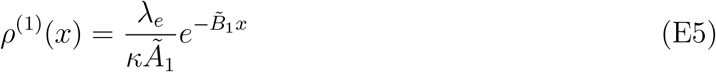

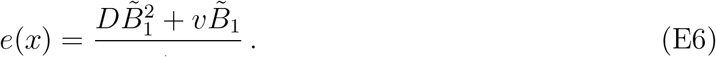

Notably the optimal enzyme density *e*(*x*) is again constant, but now depends on the drift velocity *v*.

In the region with no enzymes, *ρ*^(2)^(*x*) obeys a diffusion equation with drift, 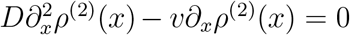, which has the solution

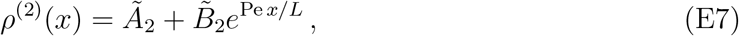

where Pe = *vL/D* is the Péclet number. As before, we evaluate the integration constants 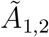, 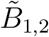 using the boundary conditions at *x* = 0 and *x* = *L*, and the continuity and smoothness conditions at *x*_0_. We again eliminate the dependence on *e_S_* by using the constraint on the total enzyme number. We are left with an expression for 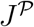 in terms of the single optimization variable *x*_0_,

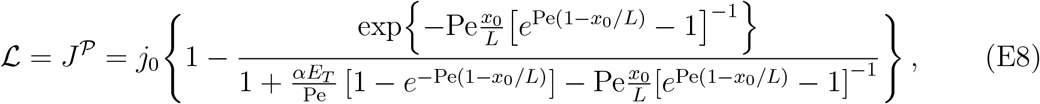

where *α* is defined as in Section B. It can be shown that Eq. E8 reduces to Eq. B9 in the limit Pe → 0.

It is not possible to find the 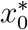 that maximizes 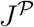 analytically. However, it is straight-forward to solve 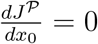 numerically for specified values of the other parameters appearing in Eq. E8. The results for the rescaled reaction flux and the rescaled net diffusive flux at the origin are shown in Fig. 3 of the main text.

## Appendix F: One-dimensional system with two sources of substrate

Here we present an example of a scenario of multiple sources of substrate with different strengths. We considered a one-dimensional system in the domain 0 ≤ *x* ≤ *L*, with reflecting boundaries at both ends. At *x* = 0 we apply a source of substrate with *j*(0) = 1, and at *x* = *L* we apply a source with *j*(*L*) = 0.3. Instead of at the boundary, loss of substrate occurs via decay with rate *σ*. We again assume linear reactions, *F*[*ρ*] = *ρ*/*K*_M_. For this system we have identified a number of different regimes of the optimal enzyme arrangement as a function of *E_T_*, shown in Fig. F.1A.

At the lowest *E_T_* values, the optimal arrangement consists of a single cluster at *x* = 0, the site of strongest substrate influx. As *E_T_* crosses a first transition level, 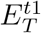, the optimal configuration features clusters of enzymes at both ends of the system: a larger one at *x* = 0 (the stronger source) and a smaller one at *x* = *L* (the weaker source). In this regime the optimal distribution of enzymes between the two poles can be calculated exactly by solving the reaction-diffusion equation with arbitrary cluster densities *e*_0_ and *e_L_*, and then optimizing over *e*_0_ and *e_L_* subject to the constraint *e*_0_ + *e_L_* = *E_T_*. The transition position 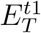 is then the value at which the resulting optimal 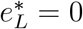. However, since the expressions are large and complex we do not include them here.

**Figure F.1.**
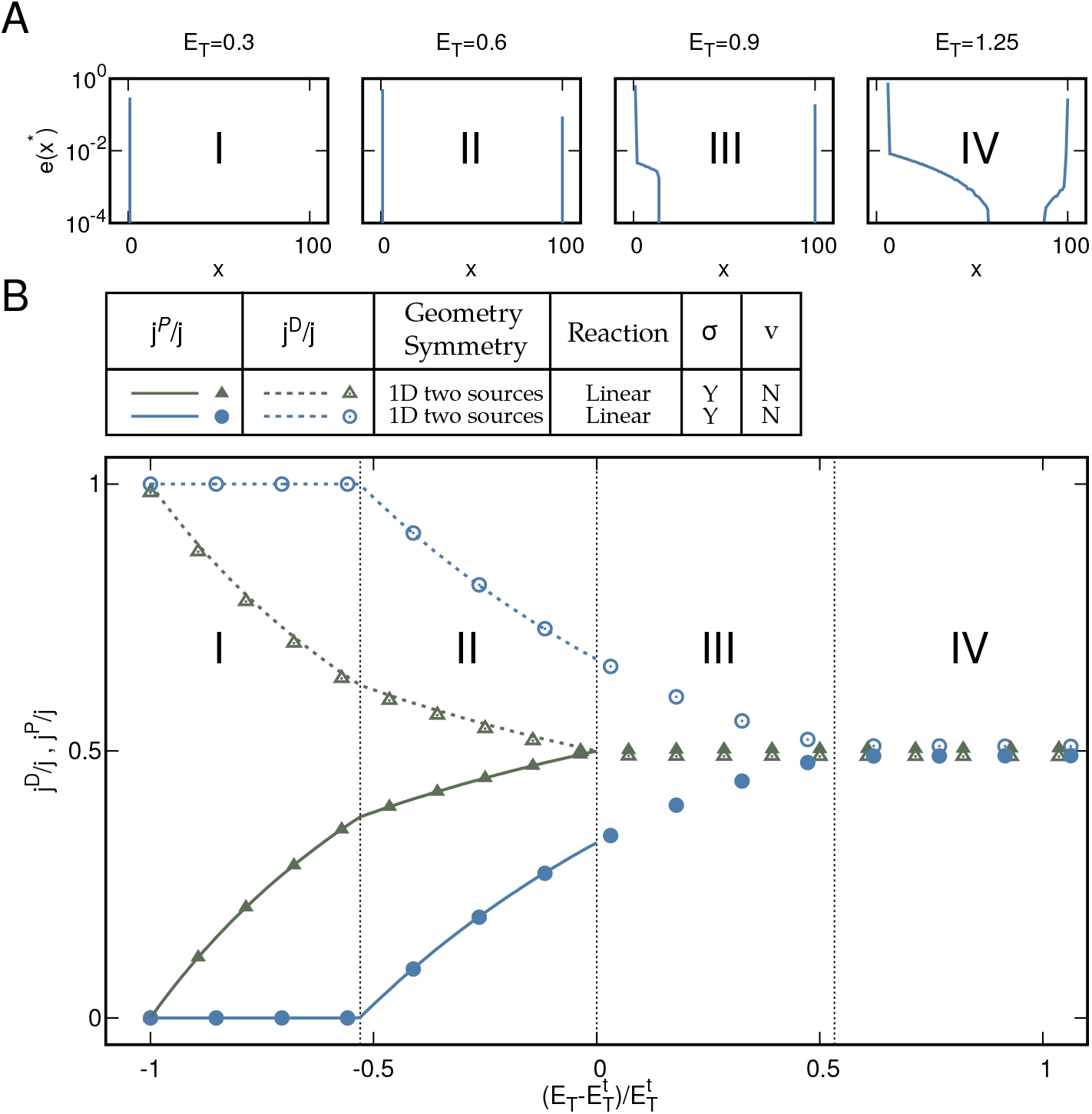
Transitions in the optimal enzyme arrangement for a one-dimensional system with two sources of substrate. The system is defined by *σ* = 1, *α* = *Lk*_cat_/(*K*_M_*D*) = 1, *v* = 0 and boundary conditions *j*(0) = 1, *k*(0) = –1, *h*(0) = 0 and *j*(*L*) = 0.3, *k*(*L*) = –1, *h*(*L*) = 0. (A) Examples of the optimal enzyme profiles for different values of *E_T_*. (B) Diffusion and reaction fluxes at *x* = 0 and *x* = *L*, normalized by the local influxes of substrate, are plotted as a function of 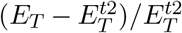. For both poles of the system, we confirm that 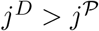 when the optimal enzyme distribution consists of only a single boundary cluster at that pole, while 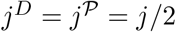 when the optimal enzyme arrangement extends a finite distance away from the pole. Lines show analytical solutions. Points show results of numerical optimization. Fig. 3 of the main text shows the same data with the fluxes at *x* = *L* plotted against 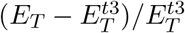.

As *E_T_* is increased further we reach a second transition at 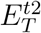, with enzymes being placed within the system in the vicinity of *x* = 0. This transition occurs when the diffusive and reaction fluxes at *x* = 0 become equal, 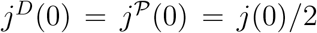 (see Fig. F.1B). At the transition value 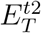 the corresponding fluxes at *x* = *L* are not equal and still satisfy the inequality condition Eq. D8 (Eq. 10 of the main text). Therefore, around *x* = *L* the optimal configuration is still purely clustered, with no enzymes at *x* < *L*. The transition threshold 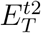 can be determined exactly by calculating the fluxes *j^D^*(0) and 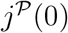 from the full solution of the system with the two optimal polar clusters.

At a larger transition value 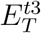, the diffusive and reactive fluxes at *x* = *L* also become equal, 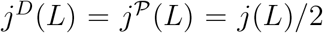 (see Fig. F.1B). For 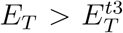, enzymes are present away from the boundaries at both ends of the system. In the regime 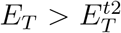 it is not possible to solve for the form of the optimal profile *e*\*(*x*), or therefore to calculate *j^D^*(*L*) and 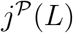. The value of 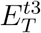 must therefore be determined numerically.

Finally, if *E_T_* is increased further, the enzyme density profiles extending from the two ends of the system will eventually coalesce, and enzymes will be present at all positions 0 ≤ *x* ≤ *L*.

It is also possible for these transitions to occur in a different order for different parameter combinations. For example, if *j*(*L*) ≪ *j*(0), the order of the first two transitions may be reversed. In this case, the condition 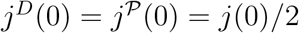 is satisfied before the transition value 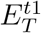 is reached. The first transition will then be from a single cluster at *x* = 0 to a regime in which there is a finite extension of enzymes from *x* = 0, but no cluster at *x* = *L*. The appearance of the second cluster at *x* = *L* may then occur subsequently, at larger *E_T_*, resulting in an optimal enzyme profile than resembles type III in Fig. F.1A.

## Appendix G: Optimal enzyme allocation algorithm for discretized systems

### 1. Derivation of the enzyme repartition rule

In this section we derive Eqs. 12, 14 of the main text, which lead to the optimal enzyme repartition rule used in step 5 of the construction algorithm. For the implementation of an optimization algorithm, we consider a system with discretized space. In this model, the densities of substrate and enzyme change from functions of position to a discrete set of variables, {*e_i_*} = {*e*(**r**_*i*_)} for *i* = 1.*N*, where *N* is the number of lattice sites. Instead of being a functional of *e*(**r**), *ρ*(**r**), the corresponding discrete Lagrangian then becomes a function of {*e_i_*}, {*ρ_i_*}, and functional derivatives 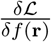 become normal derivatives 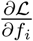.

The derivative of 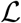 with respect to the enzyme density has an analogous form as in the continuous system,

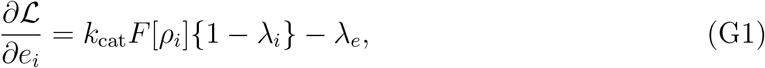

where λ_*i*_ is the Lagrange multiplier associated with the discrete reaction-diffusion equation at site *i*. We again denote the first term above by 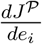, with the same interpretation as in the continuous case. This quantity is generally easily accessible in any numerical analysis: one simply increases the amount of enzymes at position *i* by a small amount, solves the resulting modified reaction-diffusion system, and calculates the resulting change in the total reaction flux. In the same way as for the continuous system, for the optimal enzyme arrangement 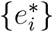 we must have that 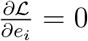 at all sites *i* where 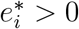, and 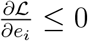 where 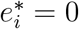.

We now consider the optimal amount of enzymes at site *i* to be a function of *E_T_*, 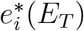. That is, there is a line **e\***(*E_T_*) in the space of **e** = (*e*_1_, *e*_2_,…,*e_N_*), that we can parameterize by the value of *E_T_*, for which the reaction flux 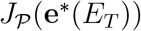 is the highest that can be achieved for that value of *E_T_*. We begin by considering the derivative of 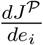 with respect to *E_T_* along the line defined by **e\***(*E_T_*), 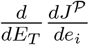. We now expand the total derivative 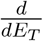 in terms of derivatives with respect to the underlying variables {*e_i_*},

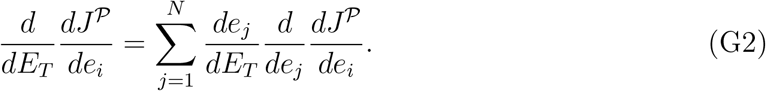

Note that here 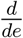 takes into account the changes in the flux both due directly to altering the enzyme density, and the resulting change in the substrate density profile {*p_i_*}. Since 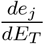 is understood to be in the direction of the line of optimal profiles, it coincides with 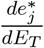.

Writing the set of *N* equations Eq. G2 in vector form we have

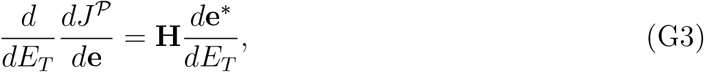

where **H** is the Hessian of 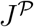, 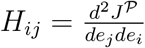.

From Eq. G1 we know that if there are 1 ≤ *n* ≤ *N* positions at which the optimal enzyme density is non-zero, then

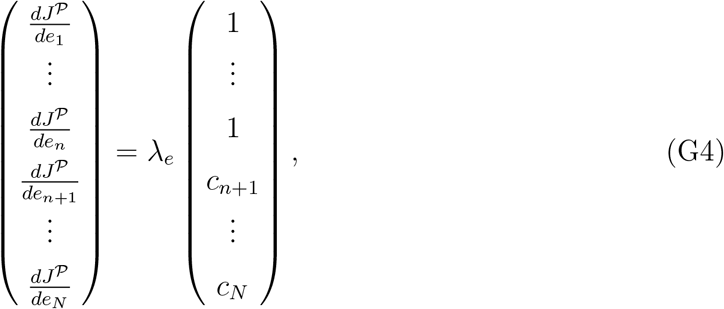

where all *c*’s lie in the range 0 < *c_i_* < 1, and we have numbered lattice sites in descending order of marginal returns. Therefore, Eq. G3 becomes

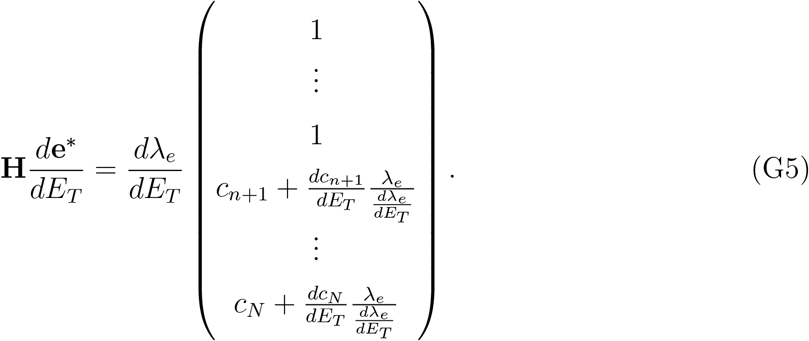

In general it is not possible to directly compute the right hand side of Eq. G5, because without knowing the direction of the optimal line it is not possible to calculate the derivatives 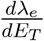 and 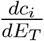. However, exploiting the fact that no enzymes should be added to the system at positions where 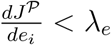, the vector 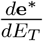 must have zeros at all positions *i* > *n*. Therefore, the *N*-dimensional system in Eq. G5 can be split into two parts. Primarily, we have the *n*-dimensional system

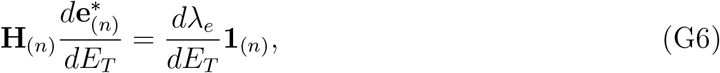

where the subscript (*n*) denotes that only the first *n* positions of the lattice are considered. Importantly, in order to solve Eq. G6 it is not necessary to know 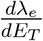 in advance. Instead, one can solve for any constant prefactor, and then subsequently use the normalization condition 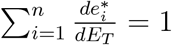 to express 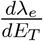 in terms of this constant. For self-consistency, the solution to Eq. G6 must also solve the residual (*N* – *n*)-dimensional system

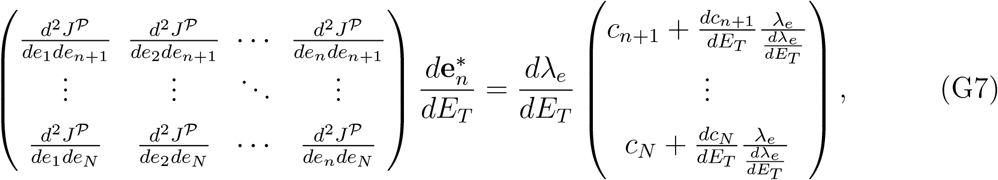

which can be interpreted as defining the evolution of the marginal returns at positions without enzymes, 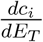.

### 2. Discretization effects

**Figure G.1.**
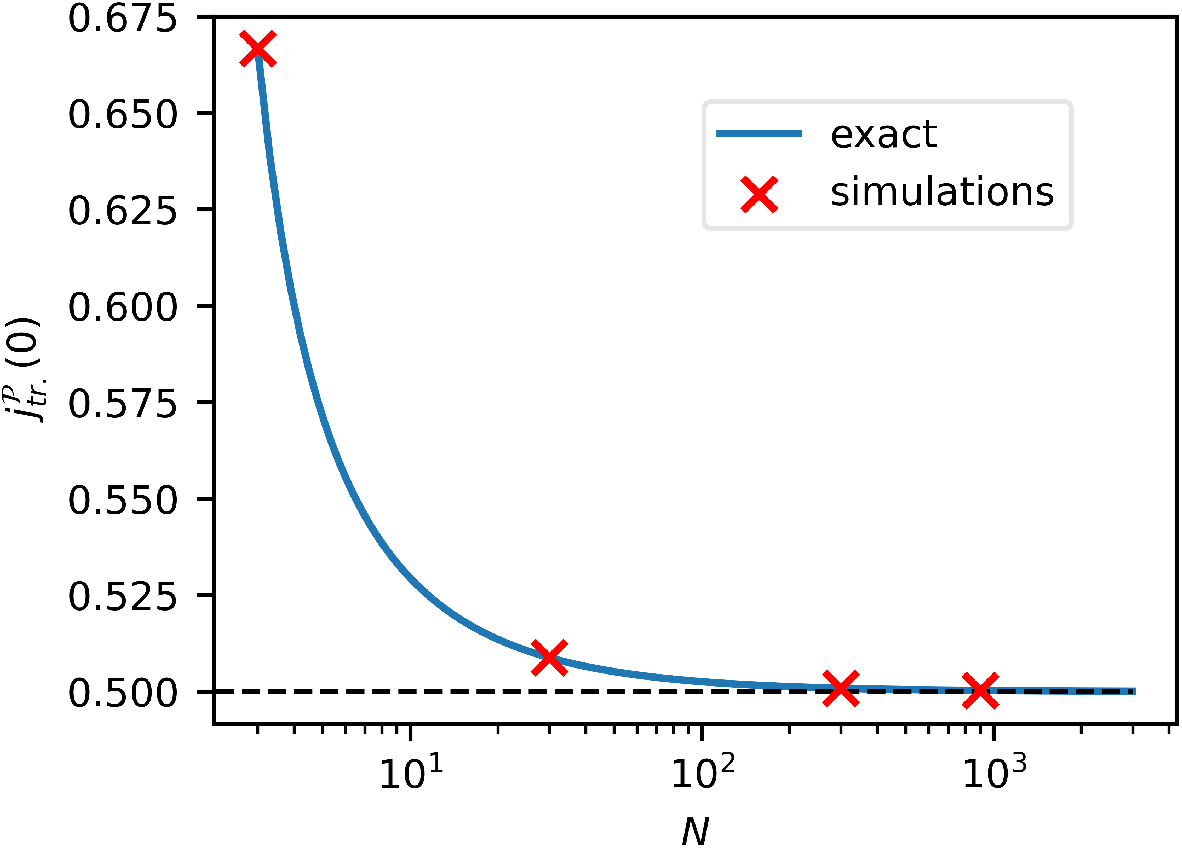
Value of the reaction flux at the transition 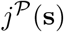 *vs* the number of lattice sites *N*. The blue line is the exact solution obtained by iteratively solving the discrete reaction-diffusion system at steady-state, while the red crosses are the result of the construction algorithm for different *N* values.

How large should the number of lattice sites *N* be to have a good approximation of the continuum limit of the system? In this section we discuss a criterion to find such a value.

In the main text we derive a general condition for the transition in the optimal enzyme arrangement from a localized arrangement into an extended profile. The condition (Eq. 10 of the main text), in combination with the boundary condition, at the transition, reads:

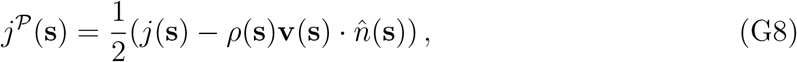

which becomes 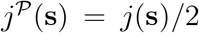 in case of no advection **v** = 0. This condition does not depend on the geometry, nor on the loss mechanism considered for the substrate and it applies locally at every position of the system boundaries with influx of substrate. As can be seen in Appendix D, the above condition is equivalent to 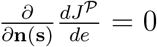 and in general is derived by combining different derivatives of the Lagrangian 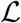. The derivatives are going to be affected by the system discretization, implying that Eq. G8 is valid only in the limit *N* → ∞.

For example in the case of the discrete version of the system described in Fig. 2 of the main text and solved in Appendix B, C, it can be shown by solving iteratively the reaction-diffusion equation, that the transition happens for 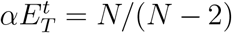 and that the reaction flux at the transition is 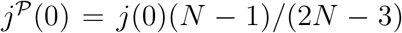. In the limit of *N* → ∞ the two limits are 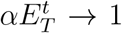 and 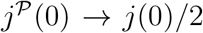, which coincide with the continuum solution, cf. Appendix B. In Fig. G.1 we can see how 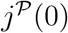 scales as a function of the number of lattice sites *N*. In blue is the exact solution, while the red crosses are the result of the construction algorithm for different discretizations. For example, we can see how for *N* = 100 the discrepancy between the continuum limit and the discrete system is approximately 0.51%.

### 3. Efficiency of the construction algorithm

In this section we compare the computational times of the construction algorithm derived in this work and a stochastic optimization algorithm that has been previously used to find the optimal enzyme arrangements in various systems belonging to the class of systems illustrated in Fig. 1 of the main text [17, 18].

The stochastic optimization algorithm takes an initial uniform enzyme configuration. Then, an iteration of the algorithm consists in taking a random amount of enzymes and moving it from a random site to a different random site. This new enzyme configuration is then the initial configuration for the next optimization step if the reaction flux generated by the new enzyme profile is higher than the previous configuration. The optimization is stopped after a maximum number of iterations of *N_iter_*. = 10^5^, or if the flux does not increase for *N_iter_*. = 10^4^. For each change in the enzyme arrangement, the steady-state reaction-diffusion equation needs to be solved. The general form of the equation for a discrete system in matrix form is (cf. Material and Methods of the main text):

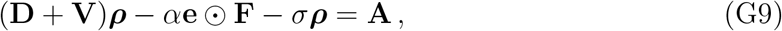

where **D** and **V** are the diffusion and advection operator on the lattice respectively, ***ρ*** the substrate density, **F** is the vector with *F_i_* = *F*[*ρ_i_*] and **A** the substrate influx vector, while ⊙ denotes the element-wise (Hadamard) product.

For the comparison of the two algorithms, we decided to consider a 1D system with two sources of substrate and linear reactions as described in Appendix F. We consider such a system because we expect the stochastic optimization algorithm to perform at its best in 1D systems with linear reactions. For such systems the diffusion matrix **D** becomes a tridiagonal symmetric matrix and Eq. G9 becomes a linear algebraic system that can be solved with the use of a tridiagonal inversion algorithm that is *o*(*N*). Since the random selection of sites is *o*(*N*^2^), we expect the stochastic optimization algorithm with a tridiagonal solver to be *o*(*N*^3^).

Our construction algorithm is also *o*(*N*^3^) since the Hessian inversion required in step 5 of the algorithm follows a Cholesky decomposition. Moreover, given the iterative nature of our construction algorithm, we have a prefactor that increases with the target amount of enzymes *E_T_* that we want to reach.

In Fig. G.2A we plot in blue circles the average computational time spent by the stochastic optimization algorithm with a solver for tridiagonal symmetric matrices over 20 repeats and with the green pluses we mark the computational time of the construction algorithm. The standard deviation of the time spent by the stochastic optimization algorithm is not shown as it is smaller than the size of the markers used. We can see that the construction algorithm is faster than the stochastic optimization with tridiagonal solver only for low amounts of *E_T_*. Passed the first transition value 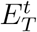, the construction algorithm gets up to 10 times slower than the stochastic optimization one with tridiagonal solver. Note, however, that the stochastic optimization algorithm returns the optimal enzyme configuration for a single *E_T_* value, whereas the construction algorithm gives the whole optimal trajectory up to the target value *E_T_*. Moreover, this is the limiting case of a system having a symmetric tridiagonal diffusion operator. In general this is not the case, e.g. higher dimensional systems. In the more general setups, one would typically use an LU-decomposition to solve Eq. G9 for linear reactions in the stochastic optimization algorithm. Such solver is *o*(*N*^3^), implying that the stochastic optimization algorithm becomes *o*(*N*^5^). In Fig. G.2A,B the computational time of the stochastic optimization algorithm using a LU-decomposition is plotted with red crosses for *N* =100 and *N* = 200 respectively. We can see how the construction algorithm out-competes the stochastic optimization one for all *E_T_* values and that the higher is the amount of lattice sites *N*, the larger is the difference in computational times as expected from the scaling argument. We therefore expects the construction algorithm to always be convenient as compared to the stochastic optimization algorithm unless the systems present simple geometries with specific symmetries. Then the construction algorithm could be slower by a factor of 10 for large *E_T_* values.

**Figure G.2.**
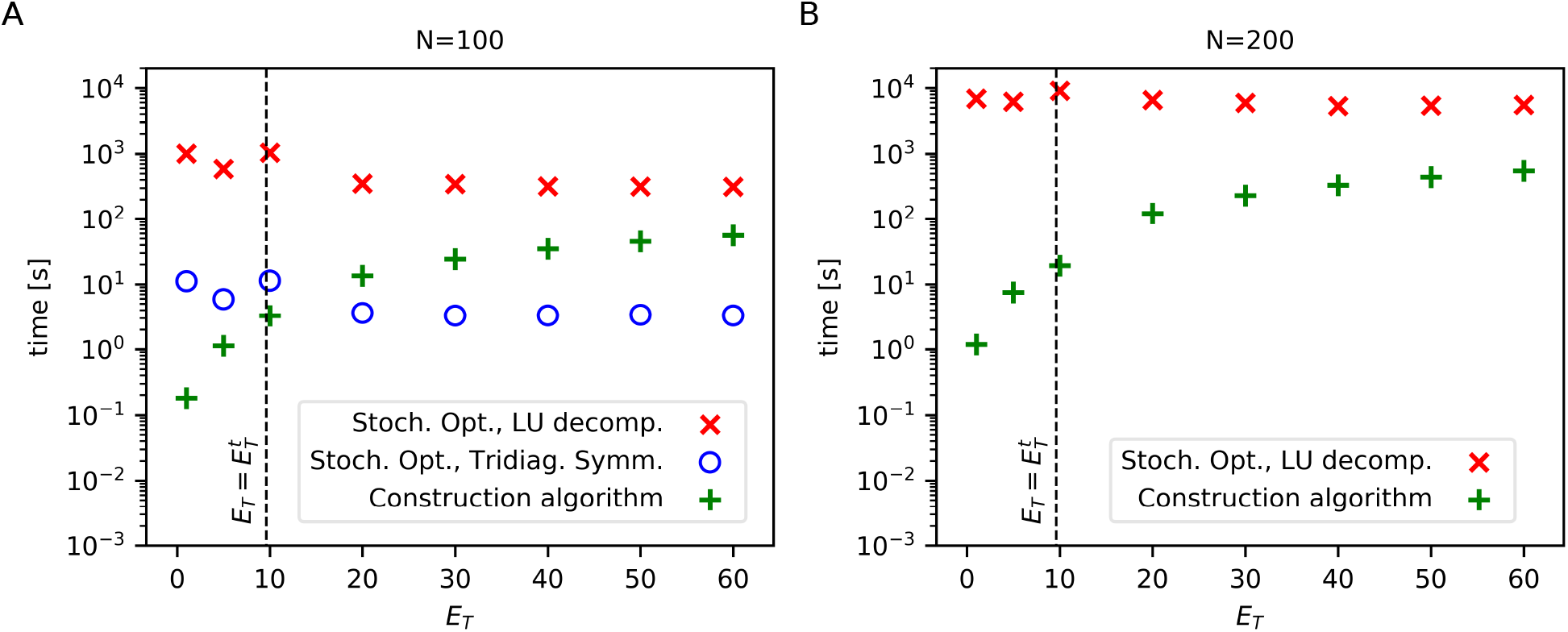
Computational time *vs* the total number of enzymes in the system for different algorithms. We mark the computational time spent by the construction algorithm derived in this work (green pluses) and the average time spent by a stochastic optimization algorithm using either a tridiagonal symmetric solver (blue circles) or an LU-decomposition algorithm (red crosses). The system under consideration is a 1D system with two sources of substrate, one for each boundary. In (A) and (B) we consider a number of lattice sites of *N* = 100 and *N* = 200 respectively. The construction algorithm out-competes the stochastic optimization algorithm with LU-decomposition, which is the standard solver used in systems with no specific symmetries and higher dimensions. The construction algorithm can be a factor of 10 slower than the stochastic optimization algorithm with a tridiagonal solver. Such a solver however can be used only for specific systems, e.g. a 1D system with a tridiagonal diffusion operator. The optimizations were conducted on a single core of a Intel(R) Xeon(R) CPU E5-2640 v3. The average time spent by the stochastic optimization is taken over 20 repeats, the standard deviation is not shown as it is smaller than the size of the markers used.

**Figure G.3.**
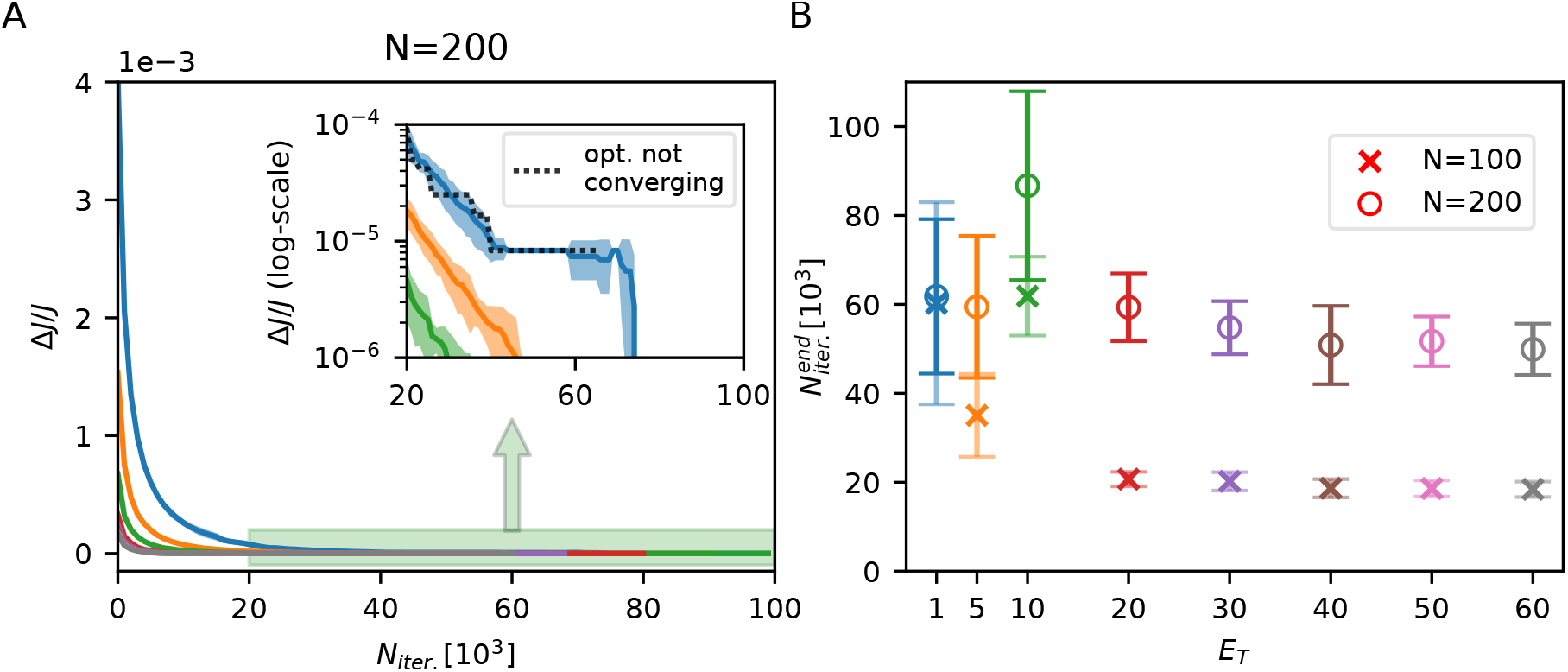
Flux comparison and convergence of the stochastic optimization algorithm vs. the construction algorithm. (A) We plot the relative difference of fluxes Δ*J/J* = 1 – *J_s,o._/J_c.a._* as a function of the number of iterations taken in the stochastic optimization algorithm for different *E_T_* values and a discretization of *N* = 200. Different colors correspond to different *E_T_* values as given by the points in panel (B). Some of the stochastic optimizations do not converge to *J_c.a._*. For example, 16 out of 20 optimization runs get stuck at Δ*J/J* 10 ^5^ for *E_T_* = 1. We plot an example of such trajectory with a black dashed line in the inset of panel (A). The optimization stops before reaching our imposed limit of *N_iter_*. = 10^5^ because the flux does not increase for 10^4^ iterations. The shaded areas correspond to the standard deviation of Δ*J/J* over 20 repeats. (B) We plot the average and the standard deviation of the number of iterations needed for the stochastic optimization to end 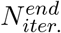. The stochastic optimization for *N* = 200 gets about 3-times higher than the number of iterations needed for *N* = 100 for *E_T_* ≥ 20. The 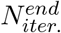 for the two discretizations get similar as *E_T_* decreases. This is due to the non-convergence of the stochastic optimization at low *E_T_* values for *N* = 200. The stochastic optimization gets stuck at sub-optimal enzyme configurations and it is stopped after 10^4^ iterations with no flux increase.

The construction optimization algorithm has also the advantage of being exact and certain to converge to the optimal solution. This is not always the case for the stochastic optimization algorithm, which may or may not converge to the global optimum. In Fig. G.3A, we plot the relative difference between the two fluxes Δ*J/J* = 1 – *J_s.o._/J_c.a._*, where *J_s.o_,* is the flux obtained via the stochastic optimization and *J_c.a_*. one of the construction algorithm. We plot Δ*J/J* for different *E_T_* values as a function of the number of iterations taken in the stochastic optimizations. None of the realizations, both for *N* = 100 and *N* = 200 resulted in negative values of Δ*J/J*, i.e. *J_s.o._* > *J_c.a_.*. In the case of *N* = 100, for any *E_T_* value and for all the repeats, Δ*J/J* goes to zero. However, in the case of *N* = 200 (Fig. G.3A), the stochastic optimization does not always converge to the flux obtained via the construction algorithm *J_c.a._*. For example, in the case of *E_T_* = 1, 16 out of 20 repeats do not converge to *J_c.a._* and they get stuck at a value of Δ*J/J* ≈ 10^-5^. We plot an example of such an optimization trajectory in the inset of Fig. G.3A (black dotted line) in log-scale. For such a trajectory, the stochastic optimization algorithm stops after *N_iter_*. ≈ 6 · 10^4^. This is better shown in Fig. G.3B, where we plot the number of iterations needed to end the stochastic optimization, 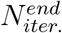, for different *E_T_* values. We can see that for a discretization of *N* = 200, 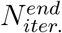 is about 3-times higher as compared to the iterations needed for *N* = 100 for *E_T_* ≦ 20.

As *E_T_* decreases, the values of 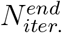 of the two discretizations get closer to each other. This is because in this regime the stochastic optimization for *N* = 200 do not always converge to the optimal enzyme arrangement. It gets stuck at some sub-optimal configuration, causing the optimization to end prematurely after 10^4^ iterations with no flux increase.

## Appendix H: Betting games with coupled outcomes and bets

### 1. Gambler repeatedly betting a fixed amount of resources

In the main text, we show that the optimal enzyme spatial allocation problem can be mapped to a betting game where a gambler bets a fixed amount of resources at every repetition of a game. In case of victory, the gambler wins an amount proportional to the bet *b_i_* on the winning event *i*, times the odds payed back to the winner *α_i_*, i.e. an amount of money *α_i_b_i_*. The long-term amount of capital won by the gambler grows arithmetically and is *C* = ∑_*i*_ *α_i_b_i_p_i_*, where *p_i_* is the probability that *i* is the winning outcome. Betting a fixed amount of resources at every repetition of the game implies that ∑_*i*_*b_i_* = *B*. We can therefore ask: what is the optimal way to repartition the bets *b_i_* under the budget constraint? To answer the question we can look at the maximum of the following Lagrangian

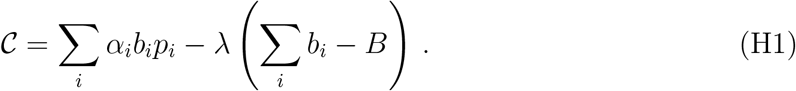

The Lagrangian is linear with respect to the bets *b_i_*. This implies that the optimal solution is to bet all the budget on the outcome that has the highest expected odds *α_i_p_i_*. Therefore in this simple scenario the gambler never diversifies the bets placed. By looking at the derivatives of the constrained capital with respect to the bets, we can see how they do not depend on any of the bets *b_i_* and are given by the sum of two terms:

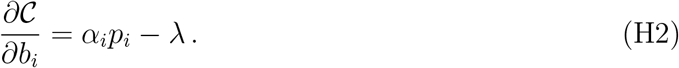

The derivative above can be interpreted as the variation of the constrained capital per extra bet placed. The first term represents the net expected gain *α_i_p_i_* per extra bet placed on the outcome *i*, if no constraint was considered. It is equivalent to the first term present in the derivative 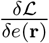, Eq. 7 of the main text, corresponding to the increase in reaction flux that would be observed upon adding extra enzymes at position **r**, if the constraints on the reaction-diffusion dynamics and the enzyme budget are neglected. The second term −λ is the marginal cost of adding extra capital and it is analogous to the – λ_*e*_ term of 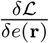.

What we are missing is an equivalent of the second term of 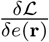, −_*V*_(**r**)*k*_cat_*F*[*ρ*(**r**)], which is generated by the coupling between *e*(**r**) and *ρ*(**r**) through the reaction-diffusion equation. In fact, in the enzyme spatial allocation problem there is a dependency between the enzymes allocated and the reaction flux generated, i.e. a dependency between the resources invested in the system and the returns obtained. We can therefore ask: what would happen to our betting problem if a feedback between the bets placed *b_i_* and the expected odds *α_i_p_i_* is introduced?

In our mapping, the bets placed on a certain outcome correspond to the enzymes allocated at a certain position, *b_i_* ↔ *e*(**r**), while the fixed budget corresponds to the total enzyme amount, *B* ↔ *E_T_*. The probability of *i* being the winning event maps to the probability of an enzyme being bound with the substrate at a certain position, *p_i_* ↔ *F*[*ρ*(**r**)], and the odds correspond to the catalytic rate, *α_i_* ↔ *k*_cat_. Since in our correspondence *α_i_* is constant, we introduce a feedback between the bets *b_i_* and the expected odds *α_i_p_i_* by considering each *p_i_* to be a function of the vector of bets *p_i_* = *f_i_*(**b**). Note, however, that a feedback could also be constructed by having the odds *α_i_* function of the bets **b**. A new functional 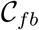 can be considered:

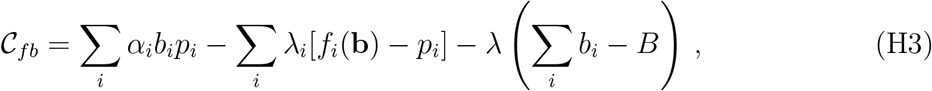

where λ_*i*_ are the Lagrange multipliers corresponding to the constraints on the probabilities *p_i_* being functions *f_i_*(·) of the bets placed **b**. The role played by the λ_*i*_ is analogous to the one played by »(**r**) in the Lagrangian 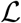 Eq. 5 of the main text. When we take derivatives of 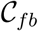 with respect to *b_i_* we now have extra terms:

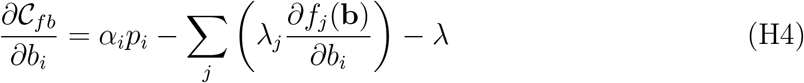

The contribution of the terms in the sum ∑_*j*_(·) comes from the coupling between the bets placed and the returns generated, analogously to the second term present in the derivative 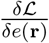, Eq. 7 of the main text. The exact form of *f_i_*(**b**) depends on the details of the model considered for the game played by the gambler.

For an optimal allocation of capital the derivative 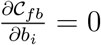, meaning that the sum of the first term with the sum ∑_*j*_(·) equals the marginal cost λ, in a similar way as what we found for the optimal allocation of enzymes by imposing 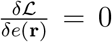. It results optimal to invest up to the point for which the net marginal gain expected from our investment (including the contribution due to the feedback) equals the marginal cost of adding extra resources. Moreover, diversification of bets can be optimal depending on the functional form of *f_i_*(·). For example, let’s consider the case of two possible winning outcomes plus a third outcome having nonzero probability of happening but zero odds. In this simple scenario and in the case of bets affecting only the correspondent outcomes, i.e. 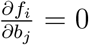 with *i* ≠ *j*, the optimal strategy is

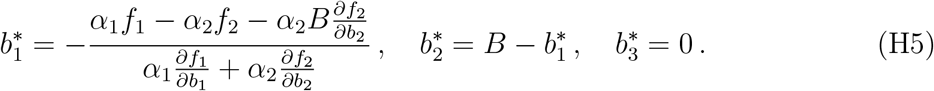

The optimal strategy results in diversified bets (i.e. 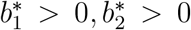), if, e.g., the events have diminishing returns (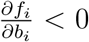, with *i* = 1, 2) and at the same time an event has higher expected odds as compared to the other 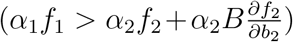. The term 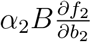 accounts for how the expected odds of the second event change due to the diminishing returns effect. Alternatively one could have diversified bets if the expected odds of an event are lower than the other 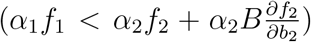 and the events present an increasing returns effect for which the probabilities of the events of being the winning ones increase with the amount of bets placed on them (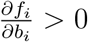, with *i* = 1,2). Moreover, if the events behave differently and for example an event presents diminishing returns and the other increasing returns, we still get optimal diversification of bets as given by Eq. H5.

To see how such optimal diversification strategies vary as a function of the total budget *B*, one can use the construction algorithm presented in the main text and derived in Appendix G. The algorithm consists in looking at the Hessian matrix of the capital 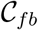 with respect to the bets placed on different outcomes, as similarly illustrated in the main text for the enzyme spatial allocation problem. The algorithm applies to any game having a linear budget constraint. This means that it can be applied to investment problems where the revenue grows arithmetically as in Eq. H3, but also problems where the revenue grows exponentially as it is the case for example for the famous Kelly problem [10]. There, instead of considering the Hessian matrix of the capital as given by 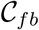, one would look at the Hessian of the capital growth rate with respect to the bets placed at different outcomes and would optimize for the growth rate as the budget gradually increases.

### 2. Gambler repeatedly reinvesting a fraction of the resources - Generalization of Kelly’s criterion

In this section we generalize Kelly’s optimal betting criterion [10] for a gambler that wants to maximize his steady state capital growth rate *G* by betting on a series of mutually exclusive events. In the original formulation of Kelly, the capital growth rate *G* has the following form:

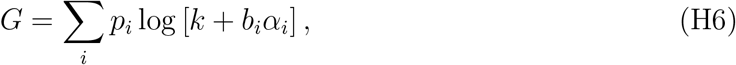

where *p_i_* is the probability for the event *i* to be the winning one, *k* is the fraction of the gambler’s capital kept, *b_i_* is the fraction of gambler’s capital invested on the event *i* and *α_i_* are the odds, i.e. the money payed back to the gambler in case *i* is the winning event.

Kelly’s aim was to determine the optimal way of partitioning the gambler’s capital given that the gambler cannot borrow nor lend money, hence his aim was to determine the optimal set of *b_i_* and *k_i_*, under the constraint ∑_*i*_ *b_i_* + *k* = 1. We can therefore consider the functional

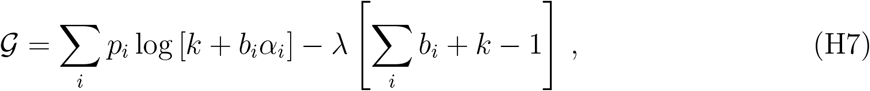

where λ is the Lagrange multiplier associated with the constraint on the gambler’s capital.

To find the optimal set of *b_i_* and *k* we perform the following derivatives:

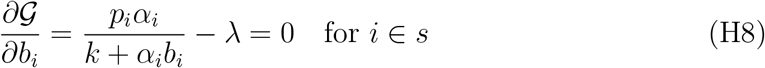

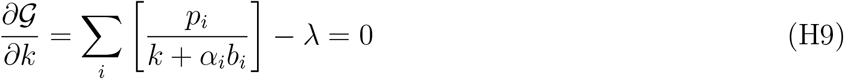

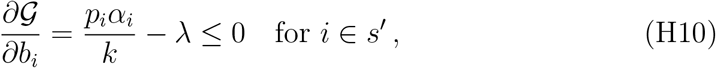

where we indicated with *s* the set of events for which it is optimal to bet and *b_i_* > 0, and with *s*′ the set of events for which betting is suboptimal and *b_i_*=0. The derivatives above can be interpreted as the variation of the constrained capital growth rate 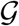 per extra bet placed H8, H10 or per extra resource kept H9. The first terms of the derivatives are the net marginal increase of *G* generated by either placing the bets H8, H10 or by keeping an extra amount of resources H9, if no constraint was considered. The −λ terms correspond to the marginal cost of adding the extra resources, analogously to the −λ_*e*_ term present in the derivative 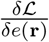, Eq. 7 of the main text. Analogously to the enzyme spatial allocation problem, we see that the optimal strategy is to bet on those events *i* ∈ *s* up to the point for which the marginal gain generated equals the marginal cost. Moreover the optimal way of repartitioning the gambler’s capital, known as Kelly criterion, can be obtained by solving for the above derivatives, together with the use of the constraint:

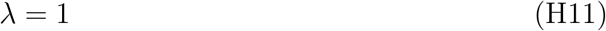

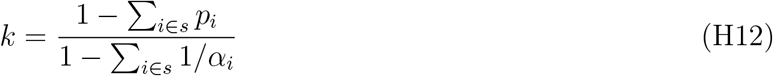

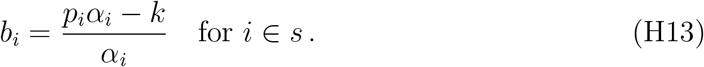

To understand how to determine the set *s* of events for which it is optimal to bet, we refer the readers to Kelly’s original paper [10], here we just say that finding the optimal *s* is similar in determining the positions at which is optimal to allocate enzymes in the class of reaction-diffusion systems considered in the main text.

Again, the variation of the constrained quantity we want to maximize per extra resource invested, i.e the derivative Eq. H8, is given by the sum of just two terms and it lacks an equivalent of the second term of 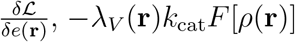, which is generated by the coupling between *e*(**r**) and *ρ*(**r**) through the reaction-diffusion equation. We can imagine to add a dependency between the probabilities of winning and the bets placed, by assuming that the probabilities *p_i_* are now functions of the bets placed *p_i_* = *f_i_*(**b**), where **b** is the vector (*b*_1_, *b*_2_,…) A new functional 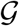 can be considered:

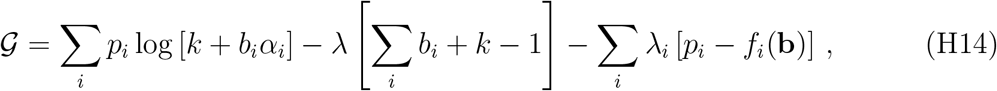

where λ_*i*_ are the Lagrange multipliers corresponding to the constraints on having the probabilities functions of the bets placed. The role played by the λ_*i*_ is analogous to the one played by λ(**r**) in the Lagrangian 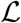 Eq. 5. When we take the derivatives of 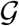 with respect to *b_i_* we now have extra terms:

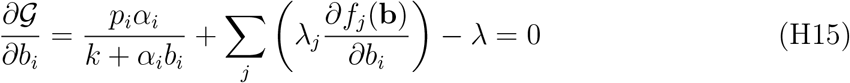

The sum ∑_*j*_(·) comes from the coupling between the bets placed and the returns generated, analogously to the second term present in the derivative 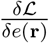, Eq. 7 of the main text. The exact form of *f_i_*(**b**) depends on the details of the model considered for the game played by the gambler.

For an optimal allocation of capital the derivative 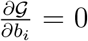, meaning that the sum of the first term with the sum ∑_*j*_(·) equal the marginal cost λ, in a similar way as what we found for the optimal allocation of enzymes by imposing 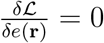.

Since the cost of adding extra resources is constant and equals −λ, we can use a similar construction algorithm as the one presented in the main text to find the optimal gambling strategy. Instead of considering the Hessian of the reaction flux with respect to the addition of enzymes at different positions, one needs to compute the Hessian of the capital growth rate with respect to the bets placed on different outcomes. Once the Hessian is found, one can derive the optimal way of repartitioning the gambler’s bet to track the optimal *G* as the total budget gradually increases in exactly the same way as we did for the enzyme spatial allocation problem.

## References

[1] K. Kumar, R. A. Mella-Herrera, and J. W. Golden, Cold Spring Harb. Perspect. Biol., 2, a000315 (2010).

[2] M. Scott, C. W. Gunderson, E. M. Mateescu, Z. Zhang, and T. Hwa, Science, 330, 1099 (2010).

[3] C. You, H. Okano, S. Hui, Z. Zhang, M. Kim, C. W. Gunderson, Y.-P. Wang, P. Lenz, D. Yan, and T. Hwa, Nature, 500, 301 (2013).

[4] A. Maitra and K. A. Dill, Proc. Natl. Acad. Sci. U.S.A., 112, 406 (2015).

[5] M. Y. Pavlov and M. Ehrenberg, Proc. Natl. Acad. Sci. U.S.A., 110, 20527 (2013).

[6] D. W. Erickson, S. J. Schink, V. Patsalo, J. R. Williamson, U. Gerland, and T. Hwa, Nature, 551, 119 (2017).

[7] D. L. Schmitt and S. An, Biochemistry, 56, 3184 (2017).

[8] F. Hinzpeter, F. Tostevin, and U. Gerland, J. R. Soc. Interface, 16, 20190444 (2019).

[9] N. Katoh and T. Ibaraki, in Handbook of Combinatorial Optimization (Kluwer Academic Publishers, 1998) pp. 159–260.

[10] J. L. Kelly, Bell Labs Tech. J., 35, 917 (1956).

[11] H. Markowitz, J. Finance, 7, 77 (1952).

[12] E. O. Thorp, in Stochastic Optimization Models in Finance (Elsevier, 1975) pp. 599–619.

[13] D. Cohen, Journal of Theoretical Biology, 12, 119 (1966), ISSN 0022-5193.

[14] P. Bauler, G. Huber, T. Leyh, and J. A. McCammon, J. Phys. Chem. Lett., 1, 1332 (2010).

[15] O. Idan and H. Hess, ACS Nano, 7, 8658 (2013).

[16] M. Castellana, M. Z. Wilson, Y. Xu, P. Joshi, I. M. Cristea, J. D. Rabinowitz, Z. Gitai, and N. S. Wingreen, Nat. Biotechnol., 32, 1011–1018 (2014).

[17] A. Buchner, F. Tostevin, and U. Gerland, Phys. Rev. Lett., 110, 208104 (2013).

[18] A. Buchner, F. Tostevin, F. Hinzpeter, and U. Gerland, J. Chem. Phys., 139, 135101 (2013).

[19] C. M. Agapakis, P. M. Boyle, and P. A. Silver, Nat. Chem. Biol., 8, 527 (2012).

[20] J. E. Dueber, G. C. Wu, G. R. Malmirchegini, T. S. Moon, C. J. Petzold, A. V. Ullal, K. L. Prather, and J. D. Keasling, Nat. Biotechnol., 27, 753 (2009).

[21] J. Zhou and B. Xu, Bioconjugate Chem., 26, 987 (2015).

[22] J. Li, Y. Kuang, J. Shi, J. Zhou, J. E. Medina, R. Zhou, D. Yuan, C. Yang, H. Wang, Z. Yang, et al., Angewandte Chemie, 127, 13505 (2015).

[23] S. Moraïs, Y. Barak, J. Caspi, Y. Hadar, R. Lamed, Y. Shoham, D. B. Wilson, and E. A. Bayer, Appl. Environ. Microbiol., 76, 3787 (2010).

[24] S.-L. Tsai, J. Oh, S. Singh, R. Chen, and W. Chen, Appl. Environ. Microbiol., 75, 6087 (2009).

[25] E. Kussell and S. Leibler, Science, 309, 2075 (2005), ISSN 0036-8075, 1095-9203.

[26] O. Rivoire and S. Leibler, J. Stat. Phys., 142, 1124 (2011), ISSN 1572-9613. >

[27] G. Fritz, N. Walker, and U. Gerland, J. Mol. Biol., 431, 4760 (2019).

[28] T. Taillefumier, A. Posfai, Y. Meir, and N. S. Wingreen, Elife, 6, e22644 (2017).

[29] O. Shoval, H. Sheftel, G. Shinar, Y. Hart, O. Ramote, A. Mayo, E. Dekel, K. Kavanagh, and U. Alon, Science, 336, 1157 (2012), ISSN 0036-8075, 1095-9203.

[30] R. E. Goldstein and J.-W. van de Meent, Interface Focus, 5, 20150030 (2015).

[31] J. F. Presley, N. B. Cole, T. A. Schroer, K. Hirschberg, K. J. Zaal, and J. Lippincott-Schwartz, Nature, 389, 81 (1997).

[32] B. Hove-Jensen, K. R. Andersen, M. Kilstrup, J. Martinussen, R. L. Switzer, and M. Willemoёs, Microbiol. Mol. Biol. Rev., 81, e00040 (2017).

[33] E. W. Miles, S. Rhee, and D. R. Davies, J. Biol. Chem., 274, 12193 (1999).

[34] J. R. Robbins, D. Monack, S. J. McCallum, A. Vegas, E. Pham, M. B. Goldberg, and J. A. Theriot, Mol. Microbiol., 41, 861 (2001).

[35] Y. E. Chen, C. Tropini, K. Jonas, C. G. Tsokos, K. C. Huang, and M. T. Laub, Proc. Natl. Acad. Sci. U.S.A., 108, 1052 (2011). >

[36] T. E. Saunders, K. Z. Pan, A. Angel, Y. Guan, J. V. Shah, M. Howard, and F. Chang, Dev. Cell, 22, 558 (2012).

[37] D. Oh, C.-H. Yu, and D. J. Needleman, Proc. Natl. Acad. Sci. U.S.A., 113, 8729 (2016).

[38] G. Hrazdina and G. J. Wagner, Arch. Biochem. Biophys., 237, 88 (1985).

[39] J. W. Graham, T. C. Williams, M. Morgan, A. R. Fernie, R. G. Ratcliffe, and L. J. Sweetlove, Plant Cell, 19, 3723 (2007).

[40] J. B. French, S. A. Jones, H. Deng, A. M. Pedley, D. Kim, C. Y. Chan, H. Hu, R. J. Pugh, H. Zhao, Y. Zhang, et al., Science, 351, 733 (2016).

[41] T. Laursen, J. Borch, C. Knudsen, K. Bavishi, F. Torta, H. J. Martens, D. Silvestro, N. S. Hatzakis, M. R. Wenk, T. R. Dafforn, et al., Science, 354, 890 (2016).

[42] M. E. Campanella, H. Chu, and P. S. Low, Proc. Natl. Acad. Sci. U.S.A., 102, 2402 (2005).

[43] H. W. Kuhn and A. W. Tucker, Proceedings of the Second Berkeley Symposium on Mathematical Statistics and Probability, 481 (1951), publisher: University of California Press.

[44] W. Karush, M. Sc. Dissertation. Dept. of Mathematics, Univ. of Chicago (1939).

[45] A. Bar-Even, E. Noor, Y. Savir, W. Liebermeister, D. Davidi, D. S. Tawfik, and R. Milo, Biochemistry, 50, 4402 (2011), ISSN 0006-2960.

[46] B. Ho, A. Baryshnikova, and G. W. Brown, Cell Syst., 6, 192 (2018).

[47] In general we do not know 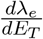 *a priori*. However, this scalar prefactor affects only the length, and not the direction, of 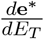. Hence it suffices to treat 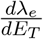 as an arbitrary constant, and subsequently to rescale the resulting 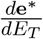 to unit length once its direction is known.

[48] L. Dinis, J. Unterberger, and D. Lacoste, EPL (Europhysics Letters), 131, 60005 (2020), ISSN 0295-5075.

[49] T. Thomik, I. Wittig, J.-y. Choe, E. Boles, and M. Oreb, Nat. Chem. Biol., 13, 1158 (2017).

[50] H. Rabeharindranto, S. Castaño-Cerezo, T. Lautier, L. F. Garcia-Alles, C. Treitz, A. Tholey, and G. Truan, Metab. Eng. Commun., 8, e00086 (2019).

[51] S.-L. Tsai, N. A. DaSilva, and W. Chen, ACS Synth. Biol., 2, 14 (2013).

[52] J.-L. Lin, J. Zhu, and I. Wheeldon, ACS Synth. Biol., 6, 1534 (2017).

[53] C. J. Delebecque, A. B. Lindner, P. A. Silver, and F. A. Aldaye, Science, 333, 470 (2011).

[54] S. Kufer, E. Puchner, H. Gumpp, T. Liedl, and H. Gaub, Science, 319, 594 (2008).

[55] K. R. Erlich, S. M. Sedlak, M. A. Jobst, L. F. Milles, and H. E. Gaub, Nanoscale, 11, 407 (2019).

[56] J. Müller and C. M. Niemeyer, Biochem. Biophys. Res. Commun., 377, 62 (2008).

[57] O. I. Wilner, Y. Weizmann, R. Gill, O. Lioubashevski, R. Freeman, and I. Willner, Nat. Nanotechnol., 4, 249 (2009).

[58] J. Fu, M. Liu, Y. Liu, N. W. Woodbury, and H. Yan, J. Am. Chem. Soc., 134, 5516 (2012).

[59] J. Fu, Y. R. Yang, A. Johnson-Buck, M. Liu, Y. Liu, N. G. Walter, N. W. Woodbury, and H. Yan, Nat. Nanotechnol., 9, 531 (2014).

